# Convergence of monosynaptic inputs from neurons in the brainstem and forebrain on parabrachial neurons that project to the paraventricular nucleus of the thalamus

**DOI:** 10.1101/2022.02.23.481069

**Authors:** Gilbert J. Kirouac, Sa Li, Shuanghong Li

## Abstract

The paraventricular nucleus of the thalamus (PVT) projects to areas of the forebrain involved in behavior. Homeostatic challenges and salient cues activate the PVT and evidence shows that the PVT regulates appetitive and aversive responses. The brainstem is a source of afferents to the PVT and the present study was done to determine if the lateral parabrachial nucleus (LPB) is a relay for inputs to the PVT. Retrograde tracing experiments with cholera toxin B (CTB) demonstrate that the LPB contains more PVT projecting neurons than other regions of the brainstem including the catecholamine cell groups. The hypothesis that the LPB is a relay for brainstem signals to the PVT was assessed using an intersectional monosynaptic rabies tracing approach. Sources of inputs included the reticular formation, periaqueductal gray (PAG), nucleus cuneiformis, superior and inferior colliculi, and the LPB. Distinctive clusters of input cells to LPB-PVT projecting neurons were also found in the dorsolateral bed nucleus of the stria terminalis (BSTDL) and the lateral central nucleus of the amygdala (CeL). Anterograde viral tracing demonstrates that LPB-PVT neurons densely innervate all of the PVT in addition to providing collateral innervation to the preoptic area, lateral hypothalamus, zona incerta and PAG but not the BSTDL and CeL. The paper discusses the anatomical evidence that suggests that the PVT is part of a network of interconnected neurons involved in arousal, homeostasis, and the regulation of behavioral states with forebrain regions potentially providing descending modulation or gating of signals relayed from the LPB to the PVT.

## Introduction

The paraventricular nucleus of the thalamus (PVT) is anatomically positioned within neural circuits that regulate behavior (Kirouac 2015) and accumulating evidence shows that this midline thalamic nucleus modulates a variety of appetitive and aversive responses (see recent reviews by Engelke et al. 2021; Barson et al. 2020; McGinty and Otis 2020; Penzo and Gao 2021; Millan et al. 2017). The connections and mechanisms involved are generating considerable interest in this rather poorly understood component of the behavioral circuits. The PVT is composed of neurons that provide dense projections via highly collateralized axons to the shell of the nucleus accumbens (NAcSh), dorsolateral bed nucleus of the stria terminalis (BSTDL), lateral central nucleus of the amygdala (CeL) (Moga et al. 1995; Li and Kirouac 2008; Parsons et al. 2007; Dong et al. 2017; Vertes and Hoover 2008; Li et al. 2021) through which the PVT has been shown to regulate a behavior (Choi et al. 2019; Choi and McNally 2017; Penzo et al. 2015; Padilla-Coreano et al. 2012; Do-Monte et al. 2017; Lafferty et al. 2020; Engelke et al. 2021; Dong et al. 2020; Pliota et al. 2018). The PVT receives afferents from the prefrontal cortex, hypothalamus and the brainstem (Li and Kirouac 2012) where these afferent inputs are processed by PVT neurons to modulate behavior via a highly divergent efferent projections (Li et al. 2021).

The PVT has been consistently identified as an area of the brain that becomes active during periods of behavioral arousal (Hsu et al. 2014; Kirouac 2015) and a number of studies have shown that the PVT modulates the hormonal and behavioral effects of both acute and chronic stress (Dong et al. 2020; Li et al. 2010; Pliota et al. 2018; Bhatnagar and Dallman 1998; Bhatnagar and Dallman 1999; Bhatnagar et al. 2003; Bhatnagar et al. 2002; Bhatnagar et al. 2000; Jaferi and Bhatnagar 2006; Jaferi et al. 2003; Heydendael et al. 2011). Homoeostatic challenges such as the physiological demands of thirst, hunger and disruptions in circadian rhythms also increase the activity of neurons in the PVT (Kirouac 2015; Colavito et al. 2015; Millan et al. 2017; Penzo and Gao 2021), suggesting that the PVT is activated by psychological and physiological stressors that have arousal effects on the CNS (reviewed in Hsu et al. 2014; Kirouac 2015). Accordingly, the PVT appears to be part of a neural network involved in coordinating a wide range of adaptive responses needed to achieve homeostasis (Li et al. 2021).

The source of brainstem afferents to the PVT has been the subject of a number of retrograde tracing studies. Early investigations with large injections of retrograde tracers in the PVT labeled neurons in a number of brainstem regions including the locus ceruleus (LC), nucleus of the solitary tract (NTS) and ventrolateral medulla (VLM) (Cornwall and Phillipson 1988; Chen and Su 1990). Studies involving more restricted injections were generally in agreement with the view that the PVT and other midline thalamic nuclei received afferents from a relatively small number of neurons in the LC, NTS and VLM (Krout et al. 2002; Krout and Loewy 2000a, b; Krout et al. 2003) some of which synthesize catecholamines (Phillipson and Bohn 1994; Otake and Ruggiero 1995; Otake et al. 1994). Brainstem catecholaminegic neurons regulate physiological and behavioral responses to homeostatic challenges via ramifying projections to many regions of the brain (Guyenet et al. 2013; Card et al. 2006; Rinaman 2011; Guyenet 2006; Dampney 1994; Aston-Jones and Cohen 2005). Recent papers have reported that catecholaminegic inputs to the PVT from the LC and VLM respectively modulates aversive stress and hypoglycemia-induced feeding responses (Beas et al. 2018; Sofia Beas et al. 2020).

However, the existence of a functionally relevant catecholaminegic input to the PVT from the lower brainstem is not unequivocally supported by all anatomical studies. For instance, our group reported that injections of the retrograde tracer cholera toxin B (CTB) restricted to the PVT rarely produced labeling in the LC whereas the NTS and VLM contained only a few scattered neurons per brain section examined (Li and Kirouac 2012). In contrast, the parabrachial nucleus (PB) located around the superior cerebellar peduncle was consistently found to contain labeled neurons with the lateral aspect of this nucleus (LPB) containing by far the most PVT projecting neurons in the lower brainstem (Li and Kirouac 2012; Kirouac et al. 2006; Li et al. 2014; Krout et al. 2002; Krout and Loewy 2000a). The LPB is densely innervated by catecholaminegic neurons in the NTS and VLM and is considered a major relay center of viscerosensory and nociceptive signals to the forebrain (Saper and Loewy 1980; Herbert et al. 1990; Moga et al. 1990; Palmiter 2018).

These anatomical observations led us to question the potential importance of a direct monosynaptic catecholamine input from the lower brainstem to the PVT and pointed to the more likely possibility that the LPB serves as an integrating and relay center for the brainstem to the PVT. The LPB also receives input from a number of forebrain regions (Tokita et al. 2009; Moga et al. 1990) and these could represent additional source of afferents integrated by LPB-projecting neurons. To address these questions, we examined the distribution of labeled neurons in brainstem catecholaminegic cell groups following injections of CTB in anterior (aPVT) and posterior aspect of the PVT (pPVT). We also tested the hypothesis that the LPB is a relay center for brain inputs to the PVT by determining the source of input cells to LPB-PVT projecting neurons using a monosynaptic rabies tracing approach called tracing the input-output organization (TRIO) method (Schwarz et al. 2015).

## Methods

### Animals

A total of 34 male Sprague–Dawley rats (University of Manitoba vivarium) were used to complete the experiments described in the paper. The rats weighed approximately 300 ± 10 g at the time of the injections and were housed on a 12:12 h light–dark cycle with food and water freely available. All experiments were carried out according to guidelines of the Canadian Council on Animal Care and approved by Research Ethics Review Board of the University of Manitoba.

### Injections of CTB for retrograde tracing experiments

Rats were anesthetized with 2–3% isoflurane and given meloxicam (2 mg/kg, s.c.) for post-surgical pain management. The animals were placed in a Stoelting stereotaxic frame and a hand drill was used to expose the brain surface above the midline thalamus. Dual injections of CTB were made using glass pipettes (approximately 40 µm diameter) directed at the aPVT (1.8 mm posterior, 1.0 mm lateral, 5.4 mm ventral, 10° angle towards the center; all coordinates are relative to bregma and bone surface) and the pPVT (3.1 mm posterior, 1.0 mm lateral, 5.3 mm ventral, 10° angle towards the center) on the right side of the brain of the same animal. The CTB injected was conjugated to Alexa Fluor-488 (AF-488-CTB, C22841, Invitrogen, Carlsbad, CA, USA) or Alexa Fluor-594 (AF-594-CTB; C22842, Invitrogen) with one of these conjugates given in either the aPVT or pPVT of the same animal. The CTB solids were dissolved at a 0.5% concentration in 0.06 M neutral phosphate buffer and 250 nl of the solution was administered over a 15 min period using a pressure injection device (Picospritzer, Park Hannifin, Hollis, NH, USA). The scalp was sutured and rats returned to their home cages for approximate 10 days prior to being deeply anesthetized with 10% chloral hydrate (600 mg/kg, i.p.) and transcardially perfused with 150 ml heparinized saline followed by 400–500 ml ice-cold 4% paraformaldehyde in 0.1 M phosphate buffer (pH 7.4). The brains were removed and post-fixed in the same fixative overnight and cryoprotected in phosphate buffered saline (PBS) containing 20% sucrose and 10% glycerin at 4 °C for 48 hrs. Coronal sections of the brain were taken at 50 µm with a cryostat (UltroPro 5000) and stored in cryoprotectant before the immunoreactions. Finally, a few rats (n = 3) were perfused and brains post-fixed as above and sections of the thalamus were obtained to carry out immunoreactions against tyrosine hydroxylase (TH) and the neuropeptide orexin-A.

### Injections of viral agents for TRIO experiments

Microinjections of viral preparations were done in two surgical procedures using similar methods as the CTB experiments (Fig. 1a). In the first procedure, rats received 350 nl injections of an adenoassociaed virus (AAV) that expresses the Cre-recombinase transgene in the retrograde direction (AAVrg-Syn1-EBFP-Cre; 7.6 x 10^12^ copies/ml; #51507-AAVrg, Addgene, Cambridge, MA, USA) in both the aPVT and pPVT using the following coordinates: aPVT (1.8 mm posterior, 1.0 mm lateral, 5.9 mm ventral, 10° angle towards the center) and the pPVT (3.1 mm posterior, 1.0 mm lateral, 5.8 mm ventral, 10° angle towards the center) in addition to a 500 nl injection of a Cre-dependent “helper AAV” (AAV2/1 hSyn-Flex-TVA-HA-G, 1.0 x 10^11^ copies/ml; NTNU Viral Vector Core, Kavli Institute, Norway) in the LPB according to the following coordinates: 7.5 mm posterior, 1.8 mm lateral, 7.0 mm ventral, 14° in the anterior to posterior direction. The helper AAV transduces the expression of TVA and a component of the rabies glycoprotein in LPB-PVT neurons that contain the Cre from injections of the AAVrg-EBFP-Cre in the PVT. The avian receptor TVA promotes infection with a rabies virus pseudotyped with the envelop protein from avian sarcoma leucosis virus type A (EnvA) and the glycoprotein (G) provides the G-deleted rabies virus the critical component for transsysnaptically infecting input cells (Lavin et al. 2019; Wickersham et al. 2007). After 3 weeks, an injection of 500 nl of G-deleted-SADB19G-EnvA-Rabies-mCherry (EnvA-RVdG-mCherry, 1.0 x 10^10^ copies/ml; NTNU Viral Vector Core) was made in the LPB using the same coordinates as for the helper AAV. Control experiments were done by excluding either the AAVrg-EBFP-Cre injections in the PVT (n =3) or the helper AAV injection in the LPB (n = 3) prior to the RVdG-mCherry injection in the LPB. Rats were perfused with fixative 7 days after the injection of the RVdG-mCherry and the brains collected for sectioning as described for the CTB experiments.

**Fig. 1.**
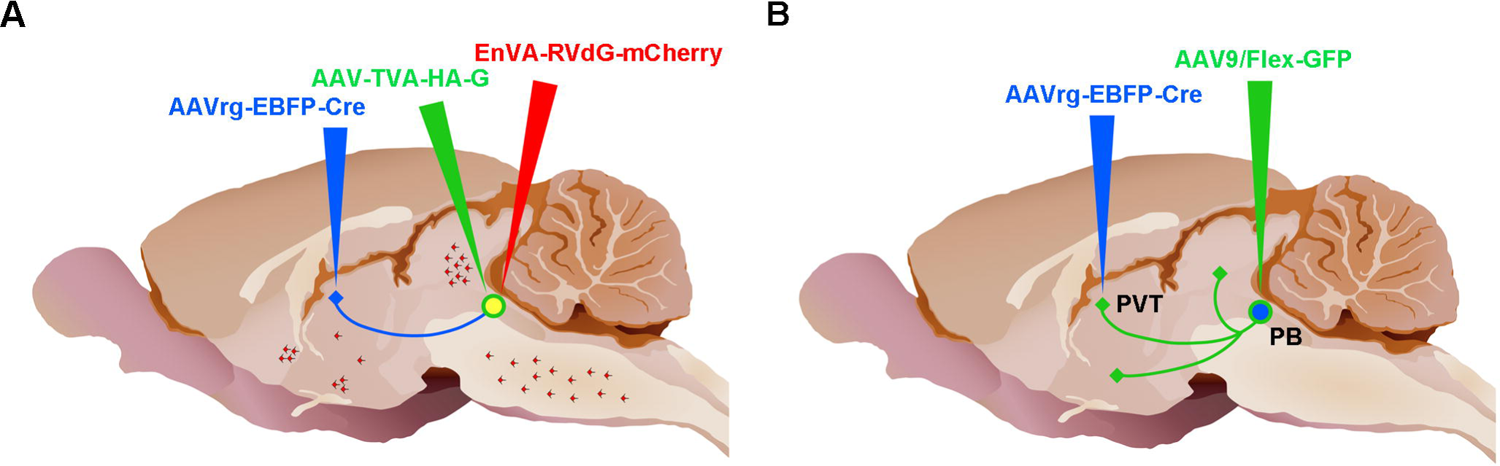
Approaches used for microinjection of viral vectors. (**a**) Injections for the TRIO experiments. (**b**) Injections for the intersectional anterograde tracing experiments. Adapted from sagittal brain images created by Gill Brown, King’s College, London. See list for abbreviations.

### Injections of viral agents for intersectional anterograde tracing experiments

The approach involves transducing Cre-recombinase in a subpopulation of projection-specific neurons using an AAV that transports in the retrograde direction (AAVrg-EBFP-Cre) in combination with an AAV that transduces neurons and fibers in the anterograde direction (Fig. 1b). This results in the tracing of axonal fiber collaterals to different regions of the brain in addition to the area initially targeted by the AAVrg-EBFP-Cre (Li et al. 2021). Rats were prepared as described above and microinjections of solutions containing an AAVrg-Syn1-EBFP-Cre preparation (500 nl) were made in the right aPVT and pPVT of the same animal followed by a microinjection of an AAV9/Flex-GFP preparation (500 nl; 5.21 x 10^13^ GC/ml, Salk Institute Viral Vector Core, La Jolla, CA, USA) in the LPB on the same side of the brain. The coordinates for the aPVT, pPVT and LPB were the same as the TRIO experiments and animals were allowed a 4-week survival period before perfusion and the removal of the brain for sectioning.

### Tissue preparation

Immunoreactions for TH were done on sections of the brainstem collected from both the CTB and the TRIO experiments. Brain sections of the LPB region were immunoreacted for human hemagglutinin (HA) to visualize the neurons transduced by the helper virus in the TRIO experiments. Finally, every sixth section was reacted for neuronal nuclear protein (NeuN) in some cases for each group of experiments to help establish the neuronal labeling according to cytoarchitectural boundaries of the rat brain. All reactions were done in free-floating sections at room temperature and involved placing sections in a blocking solution of PBS containing 5% donkey serum, 0.3% of Triton X-100, and 0.1% of sodium azide for 1 hr prior to incubation in the antiserum. The primary and secondary antibodies were diluted in the blocking solution and sections were rinsed in PBS between each step. The immunoreactions were done as described below followed by a final set of rinses before mounting and placing a coverslip using Fluromount-G (Southern Biotech, Birmingham, AL, USA).

### Tyrosine hydroxylase (TH)

Sections of the brainstem that contained CTB labeled neurons or mCherry neurons were incubated in a solution containing a mouse anti-TH monoclonal antibody (1:10,000; T-2928, Sigma-Aldrich, Oakville, ON, Canada) for 2 days. After several rinses, sections were transferred to a secondary antibody solution containing Alexa-Fluor Plus 647 (1:3000, A32787, Invitrogen) for 2 hrs. Sections of PVT were incubated in the same TH antibody solutions before transferred to a secondary Alexa-Fluor Plus 594 donkey anti-mouse antibody (1:3000; A21203, Invitrogen) for 2 hrs.

### Orexin-A

sections were incubated in a solution containing a rabbit anti-orexin A antibody (1:1000; AB3704, Millipore, Temecula, CA, USA) for two days followed by incubation in a solution containing Alexa-Fluor Plus 594 donkey anti-rabbit antibody (1:3000; A21207, Invitrogen) for 2 hrs.

### Human hemagglutinin (HA)

After one hour in the blocking solution, every sixth section of the LPB was incubated in a primary antibody solution containing rabbit anti HA-Tag (C29F4) monoclonal antibody (1:1000; 3724s, Cell Signaling Technologies, Danvers, MA, USA) overnight followed by incubation in a solution containing a secondary Alexa-Fluor Plus 488 donkey anti-rabbit antibody (1:3000; A21206, Invitrogen) for 2 hrs.

### Neuronal nuclear protein (NeuN)

Sections were incubated in a primary mouse anti-NeuN antibody (1:1000; MAB377, Chemicon, Temecula, CA, USA) overnight followed by incubation in a solution containing Alexa-Fluor Plus 488 donkey anti-mouse (1:3000; A21202, Invitrogen) for 2 hrs.

### Image and quantitative analysis

Images showing the distribution of neurons and fibers were produced from compiled stacks of frames captured using Zeiss Axio Observer Z1 microscope equipped with Axiocam 503 mono camera. The frames were taken with 10x objective lens with X and Y and Z (2 µm increments) stacks set to cover a region of interest. The exposure time was adjusted for each individual channel to optimize the images captured and kept consistent for each case in the different experiments. Images were processed using “Stitch” to fuse the X and Y tiles and then “Extended Depth of Focus” for the Z stacks (Zen Blue, Zeiss) to produce the tiled images.

Mapping and quantification of labeled neurons in the various experiments were done using a standardized gamma setting for each color channel. The contrast of the composite images was adjusted in Adobe Photoshop to produce the final images for the figures. Cell counts and mapping of the labeled neurons were done according to methods described below.

### Retrograde CTB tracing experiments

Single-labeled neurons were identified and marked on images viewed in Zen Blue with the appropriate color channel whereas multi-labeled neurons were identified in images with color channels merged. The soundness of single-cell multi-labeling was further evaluated by switching between color channels to confirm that a clear overlap of the signal was within the same soma. Images with marked single- and multi-labeled neurons were imported to Adobe Illustrator CS4 and overlaid with an image file of the appropriate anatomical level of the brain from the digital atlas of the rat brain (Paxinos and Watson 2009). The boundaries of brain nuclei and fiber bundles from the atlas images were adjusted slightly to correspond to the microscopic image. The merged image was exported to Adobe Photoshop and the locations of the labeled neurons in the brain were generated by manually placing a distinctive symbol to indicate the location of a labeled neuron. The final image files of individual stereotaxic levels were used to generate the figures displaying labeled neurons on selected stereotaxic levels of the brain.

### TRIO experimenst

The number of starter cells near the superior cerebellar peduncle and input cells for the entire brain were quantified on coronal sections captured at 300 µm intervals. Starter cells were identified as neurons with unambiguous co-distribution of mCherry and HA within the same neuronal cell body using the same methods as the CTB experiments. The number of RVdG-mCherry input cells for the different regions of the brain was quantified for those regions that had 50 or more input cells over the series of sections. The mCherry labeled cells were marked using a circular symbol on the captured images in Photoshop and the number of marked cells was quantified using ImageJ (Fuji).

### Intersectional anterograde tracing experiments

The number of GFP labeled neurons in the PB was quantified by counting the single-labeled GFP neurons from sections spanning the nucleus. Figures showing the labeling were produced from images captured as described above.

## Statistical Analyses

The data for the CTB tracing experiments were analyzed using one-way ANOVA followed by Bonferroni multiple comparison tests using the statistical software OriginPro 8 (OriginLab Corporation, Northampton, MA, USA). An adjusted value of p < 0.05 was considered to be significant and the data are presented as mean ± SEM.

## Results

### CTB retrograde tracing

It is evident that TH fiber labeling in the aPVT and pPVT is weak compared to orexin-A fiber labeling (Fig 2a). Retrograde tracing experiments were done to further evaluate the extent that neurons in the lower brainstem contribute to TH fibers in the PVT. Four cases with combination injections of CTB restricted to the aPVT and pPVT were used to generate the figures and data analysis (Fig. 2b). Table 1 displays the results of the analysis of the number of CTB neurons in the lower brainstem areas containing the catecholamine cell groups and the LPB. The LPB contained the highest number of neurons from injections of CTB in the aPVT and pPVT whereas the medial PB (MPB) contained few neurons (Figs. 3a, b, 4a, b). The brainstem regions containing catecholamine cell groups contained a relatively small number of retrograde labeled neurons (Figs. 3, 4). Table 2 displays the number of CTB retrograde labeled neurons that are TH positive indicating that catecholamine neurons represent a small fraction of the neurons in the pons and medulla that project to the PVT. Indeed, only a few neurons per section were observed in the catecholamine cell groups of the VLM (Figs. 3e-g, 4c) and the NTS (Figs. 3e-g, Figs 4d); none in the LC (Figs. 3a-c, 4a, b), and a small number of cells in the subceruleus region (Figs. 3a, b, 4b), a ventral extension of the A6 group (Bucci et al. 2017). In summary, the CTB tracing results support our previous observations that brainstem neurons innervating the PVT originate primarily from the LPB and not from the LC, NTS, VLM or other areas of the lower brainstem (Li and Kirouac 2012; Kirouac et al. 2006).

**Fig. 2.**
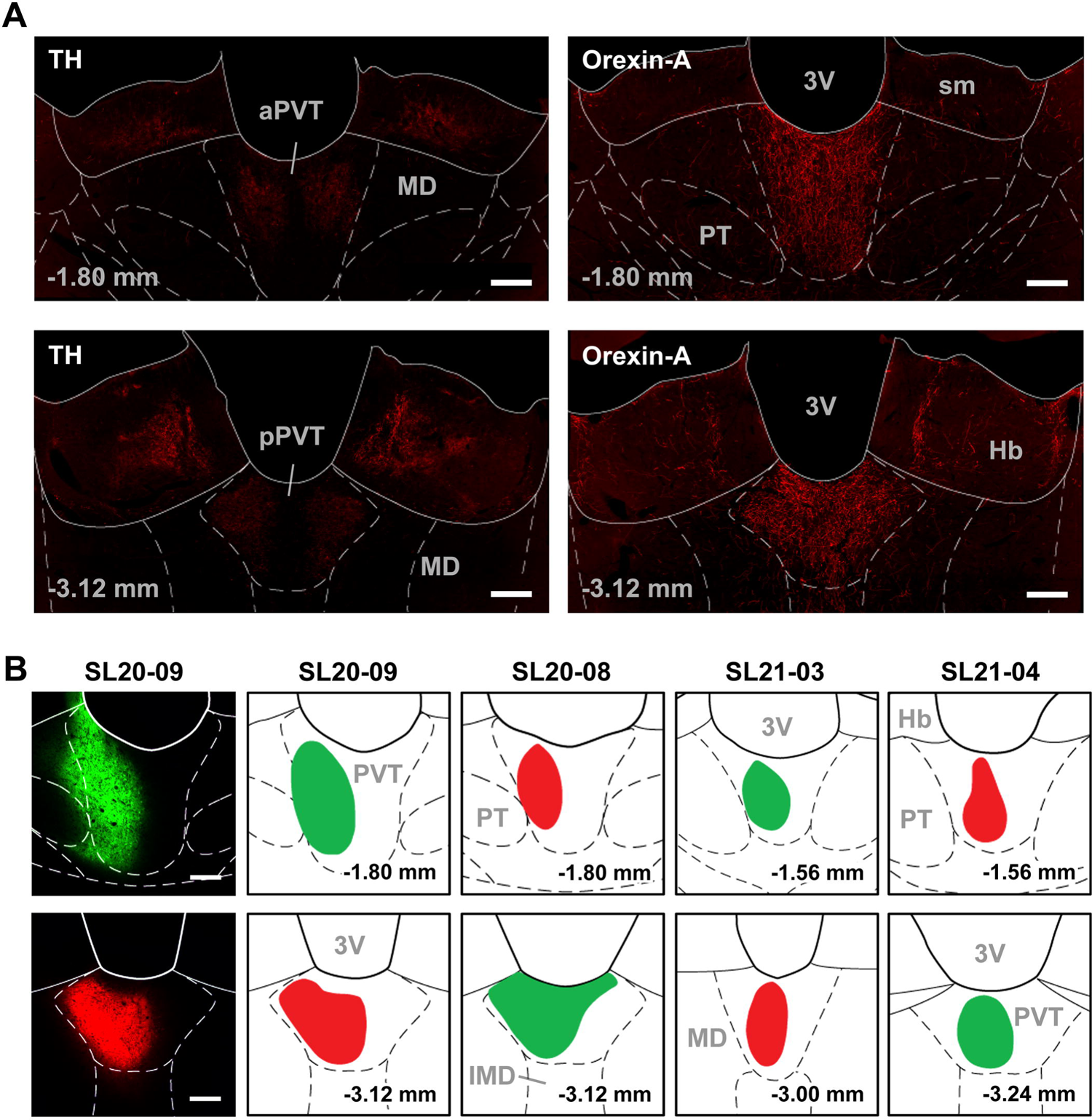
Tyrosine hydroxylase (TH) and orexin-A fiber labeling in the PVT and the location of the CTB injections in the PVT. (**a**) Immunofluorescence for TH is comparatively weaker (left column) than that for orexin-A (right column) in the aPVT (top row) and pPVT (lower row). (**b**) Location of combination injections of AF-488-CTB (green) and AF-594-CTB (red) in the aPVT (upper row) and pPVT (lower row) in 4 cases (columns) used for the retrograde tracing analysis. See list for abbreviations. Numbers at the bottom represent distance from the bregma. Scale bars: 200 µm.

**Fig. 3.**
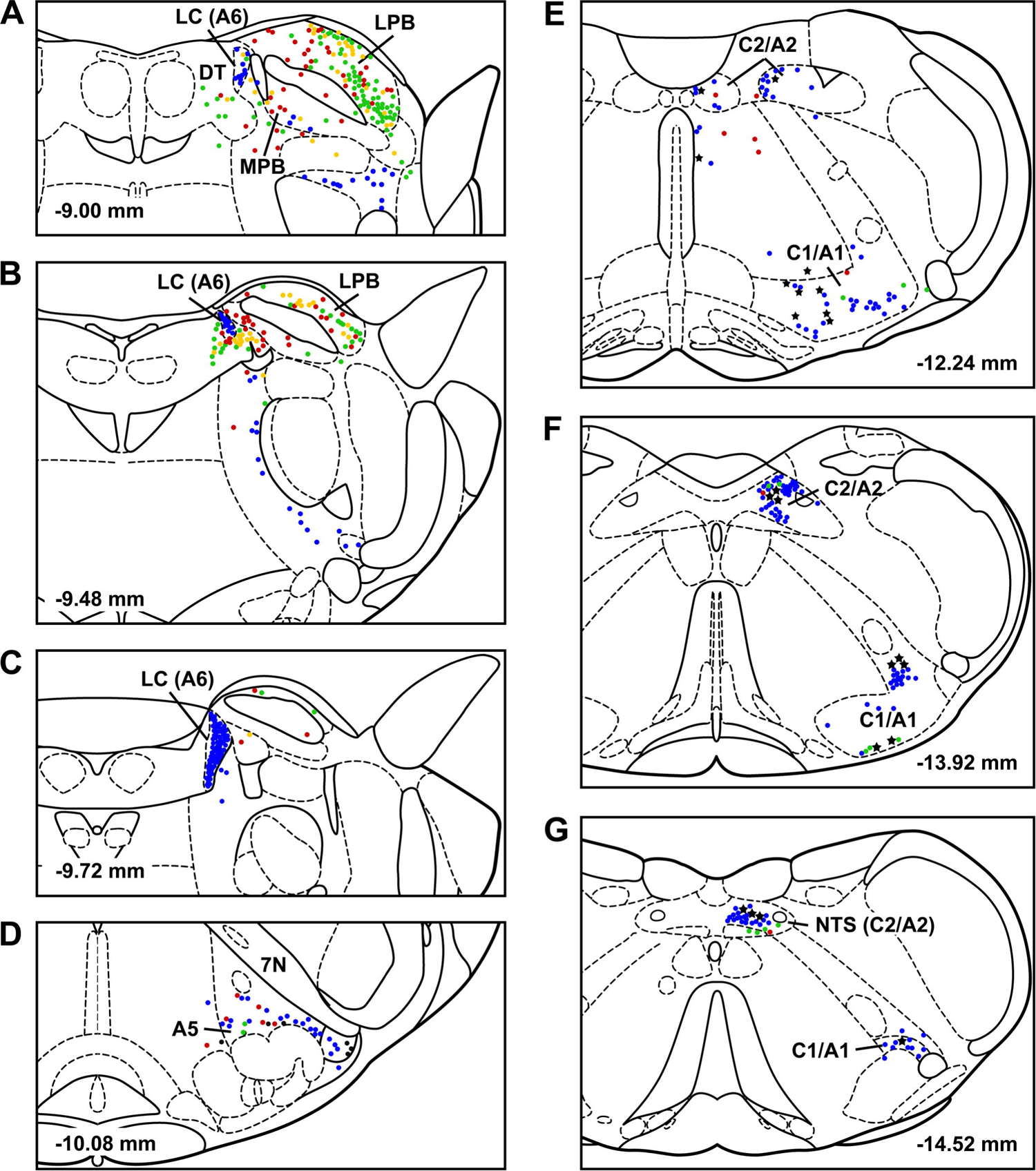
Location of labeled cells in the lower brainstem following injections of CTB in the PVT. Single CTB labeled neurons from injections in the aPVT (red) and pPVT (green) and cells projecting to both the aPVT and pPVT (yellow) are shown along with the location of TH positive cells (blue) as well as cells labeled with both CTB from either the aPVT or pPVT and TH (black stars) for case SL20-09 plotted onto a series of drawings of a rat stereotaxic atlas (Paxinos and Watson 2009) extending from the pons to the caudal medulla (**a-g**). See list for abbreviations and the numbers at the bottom represent distance from the bregma.

**Fig. 4.**
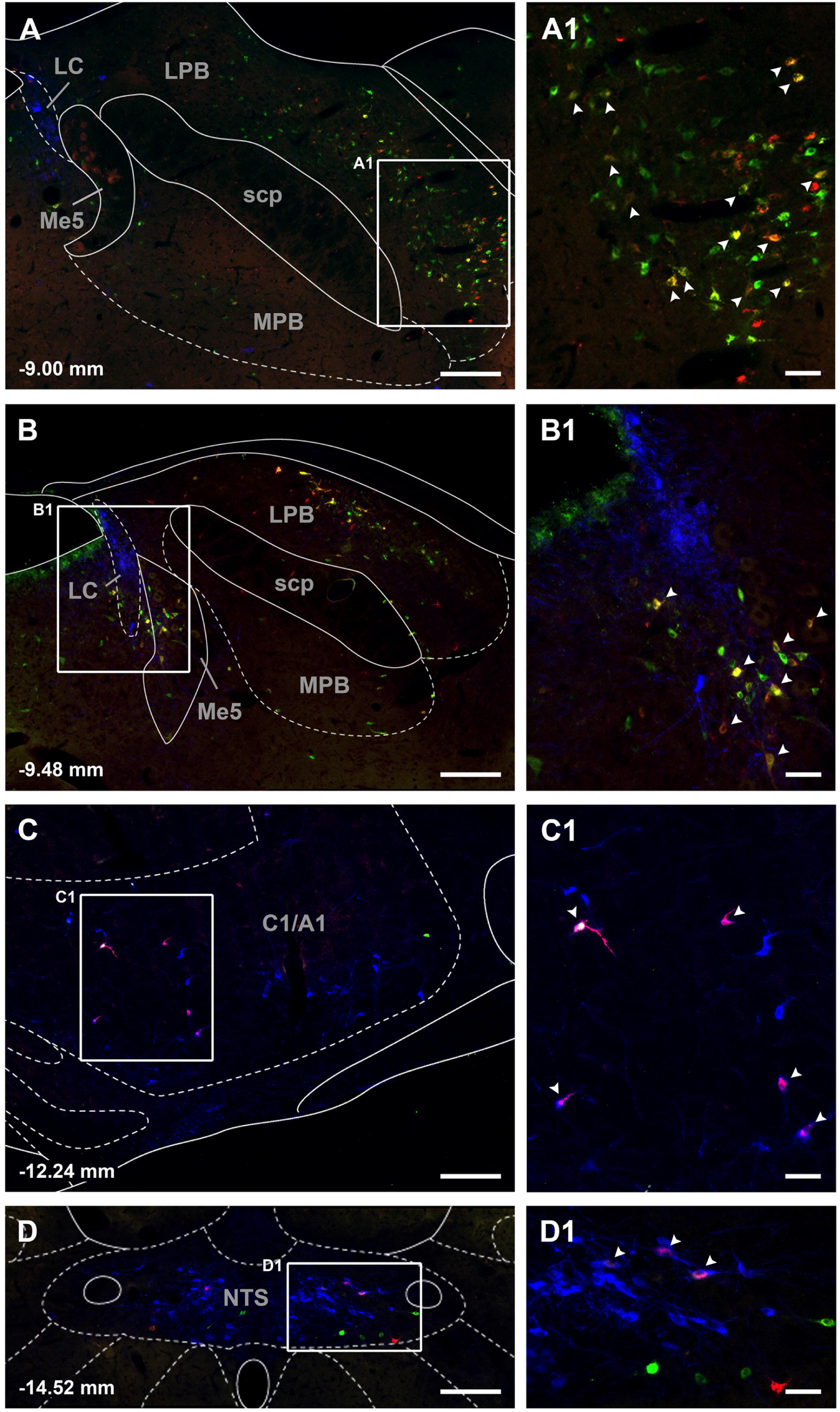
Images of CTB labeled neurons in the pons and medulla. Single CTB labeled neurons from injections in the aPVT (red) and pPVT (green) in addition to cells projecting to both the aPVT and pPVT (yellow-orange, arrow heads in **a1** and **b1**) are visible for case SL20-09. Also visible are single labeled TH positive cells (blue) and double labeled TH and CTB cells (cyan) as indicated by arrow heads in the numbered insets (**c1** and **d1**). The images are arranged from anterior to posterior levels (**a-d**) with the numbers at the bottom representing distance from the bregma. See list for abbreviations. Scale bars: 200 µm for the images, 50 µm for the insets.

**Table 1.**
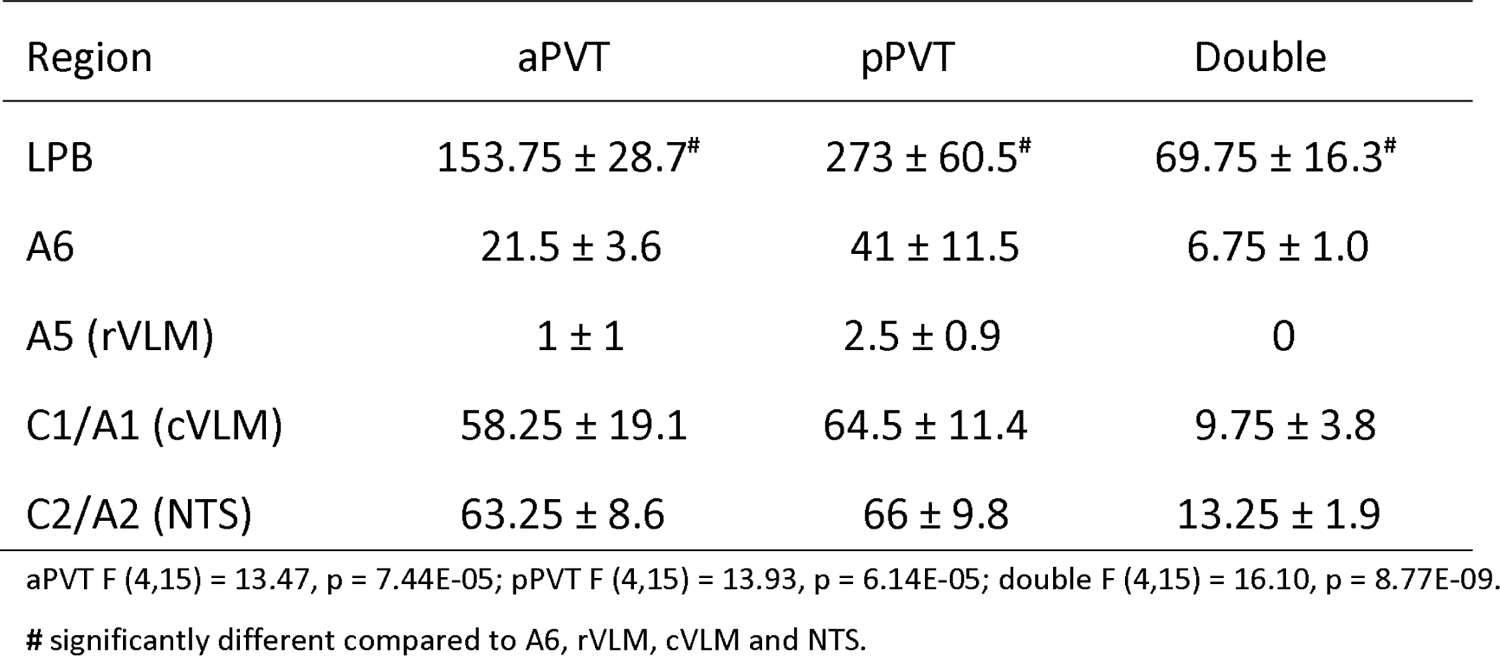
Number of retrograde CTB cells according to brainstem regions

| Region | aPVT | pPVT | Double |
| --- | --- | --- | --- |
| LPB | 153.75 ± 28.7<sup>#</sup> | 273 ± 60.5<sup>#</sup> | 69.75 ± 16.3<sup>#</sup> |
| A6 | 21.5 ± 3.6 | 41 ± 11.5 | 6.75 ± 1.0 |
| A5 (rVLM) | 1 ± 1 | 2.5 ± 0.9 | 0 |
| C1/A1 (cVLM) | 58.25 ± 19.1 | 64.5 ± 11.4 | 9.75 ± 3.8 |
| C2/A2 (NTS) | 63.25 ± 8.6 | 66 ± 9.8 | 13.25 ± 1.9 |

aPVT F (4,15) = 13.47, p = 7.44E-05; pPVT F (4,15) = 13.93, p = 6.14E-05; double F (4,15) = 16.10, p = 8.77E-09.
<sup>#</sup> significantly different compared to A6, rVLM, cVLM and NTS.

**Table 2.** Number of CTB and TH positive cells in the brainstem

| Region | aPVT | pPVT | Double |
| --- | --- | --- | --- |
| C1/A1 (cVLM) | 22.5 ± 6.0* | 29.25 ± 8.5* | 15.5 ± 5.0* |
| C2/A2 (NTS) | 12.5 ± 1.9 | 22.5 ± 6.7 | 7.5 ± 2.4 |
| A6 | 0.25 ± 0.25 | 0.25 ± 0.25 | 0.25 ± 0.25 |
| A5 (rVLM) | 0 | 0 | 0 |

aPVT $F(3,12) = 11.71$ , $p = 7.06E-04$ ; pPVT $F(3,12) = 7.78$ , $p = 3.78E-03$ ; double $(3, 12) = 7.04$ , $p = 5.5E-03$ .
\*significantly different compared to A6 and rVLM; double refers to CTB+/TH+ cells from the aPVT and pPVT

### TRIO experiments

Starter cells were identified by the presence of mCherry and HA in the same PB neuron (Fig. 5). The number of input cell per PB-PVT starter cell was quantified for the whole brain in three cases where the AAVrg-Cre injections were limited to the aPVT and pPVT (based on the presence of the reporter EBFP, see intersectional anterograde tracing results for examples).

**Fig. 5.**
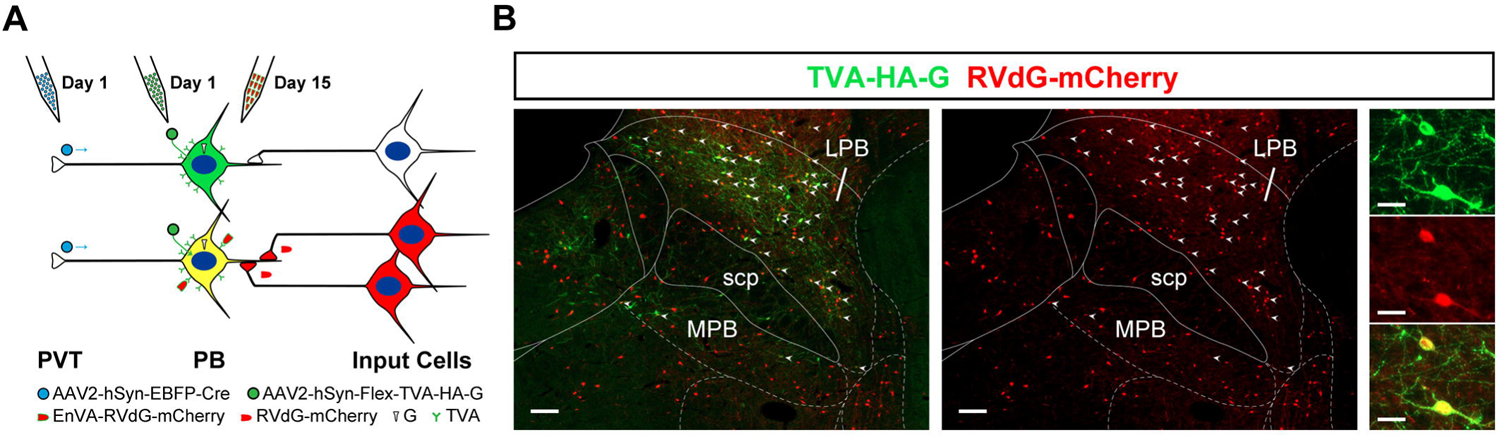
Schematic of the TRIO approach (**a**) and resulting starter cells in PB neurons that project to the PVT (**b**). The presence of starter cells is indicated by small white arrowheads at low magnification (left side) and examples are shown at higher magnification (right side) in (**b**). See list for abbreviations. Scale bars: 50 µm for the lower magnification images and 10 µm for the higher ones.

Starter cells were located primarily in the LPB (Figs. 5b, 6) in a pattern that is similar to what was observed in the CTB tracing experiments. A small number of starter cells were located in the MPB and in the transition area between the LPB and the nucleus cuneiformis immediately anterior to the LPB. The number of starter cells was consistent for the 3 cases examined (221, 209, and 205 cells in the PB sections counted).

**Fig. 6.**
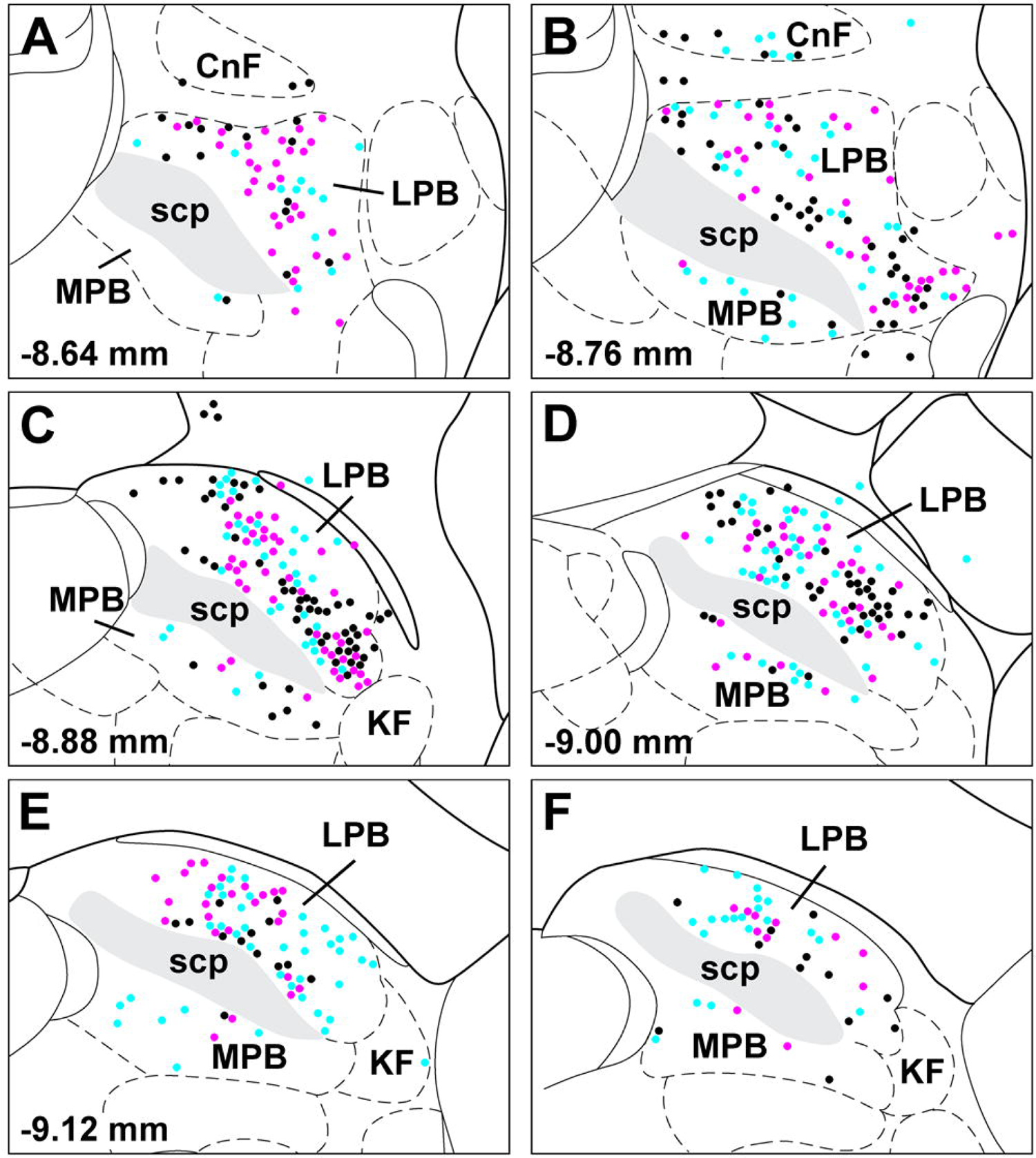
Location of starter cells in the cases used for quantitative analysis of input cells to the LPB-PVT neurons. The location of the starter cells for each case (n = 3) is uniquely color coded and is shown from from anterior to posterior levels of the PB (**a-f**). See list for abbreviations. Numbers at the bottom represent distance from the bregma.

Figure 7 shows the number of input cell per starter cell from regions of the brain that consistently contained cells in the 3 cases quantified. There was a significant difference in the number of the input cells provided in the different regional groups quantified (F_(20,42)_ = 10.63, p < 0.0001) with the reticular formation (RF) of the medulla being significantly greater than a number of other brain regions (Fig. 7). The midbrain and pontine RF also provided a significantly greater numbers of input cells in brain regions not exclusively considered as the RF (Fig. 7). Input cells were broadly scattered in the RF of the midbrain (Figs. 8a-d, 9a-c), pons (Figs, 8e-f, 9d) and medulla (Figs. 8g-h, 9e-f). The lateral substantia nigra including the transitional region between this region and the retrorubral field embed in midbrain RF were a consistent source of input cells (Figs. 8a-c, 9a-b) but these input cells were not immunopositive for TH despite being intermixed with dopaminergic neurons (data not shown). Subregions of the periaqueductal gray (PAG) including the dorsal, lateral and ventrolateral regions were a regular source of input cells to LPB-PVT neurons (Figs. 8a-d, 9a-c). Both the superior and inferior colliculi of the tectum provided a substantial number of input cell (Figs 8a-f, 9a-d) as did the nucleus cuneiformis (CnF) and the medial and lateral PB (Fig. 8e-f, 9c-d). Input cells were scattered in the NTS and VLM over a number of stereotaxic levels (Figs. 8g-h, 9e-f), most of which were immunonegative for TH (data not shown). A notable collection of input cells was observed in the anterior NTS and the adjacent parvocellular reticular nucleus in medulla, a distinctive nucleus of the medullary RF (Figs. 8g, 9e).

**Fig. 7.**
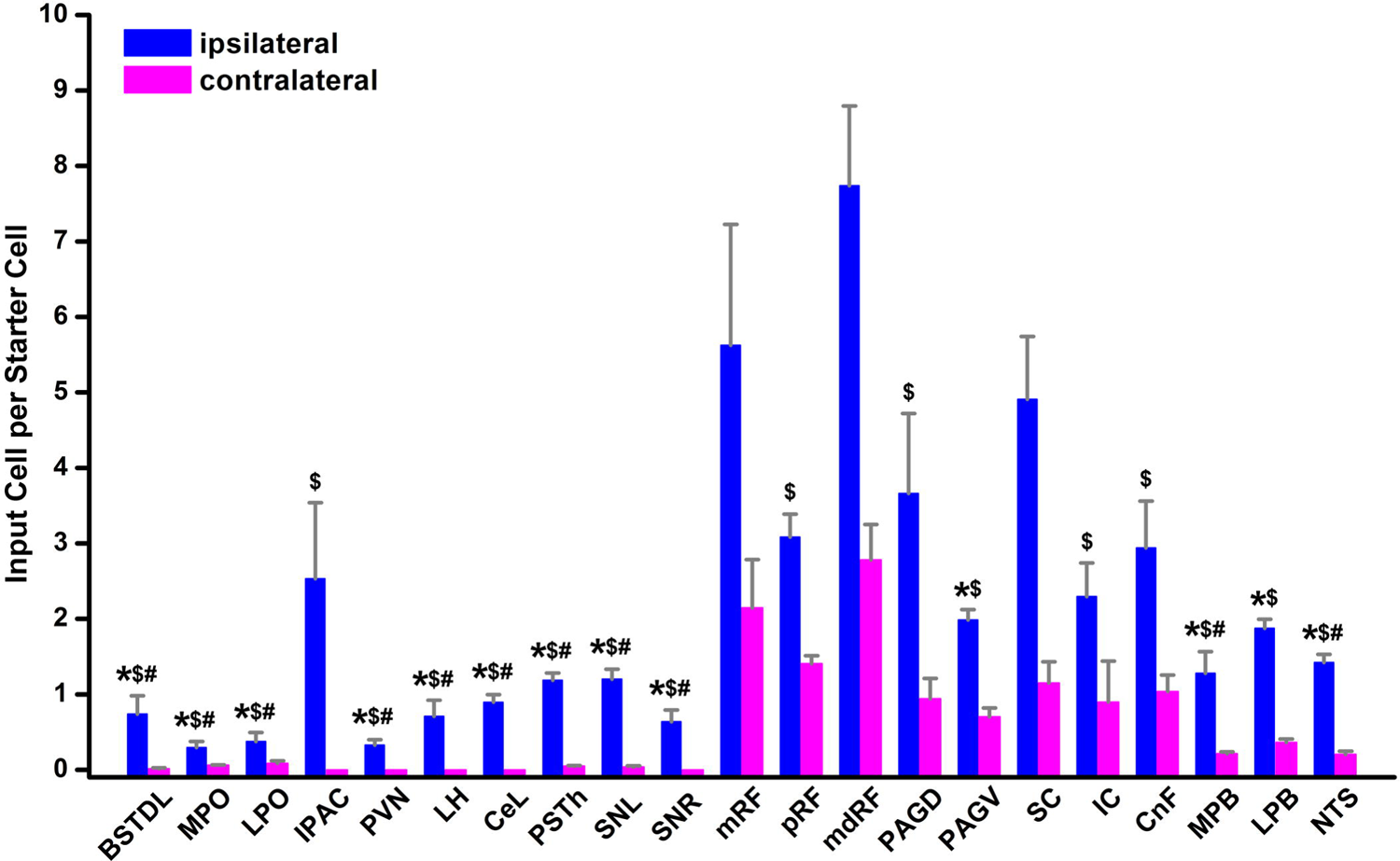
Number of input cell per starter cell from different regions of the brain making synapses on LPB-PVT projecting neurons. The brain regions quantified are listed from anterior to posterior levels. See list for abbreviations. *significantly different from mRF, ^$^ significantly different from mdRF, **^#^** significantly different from SC; p < 0.05.

**Fig 8.**
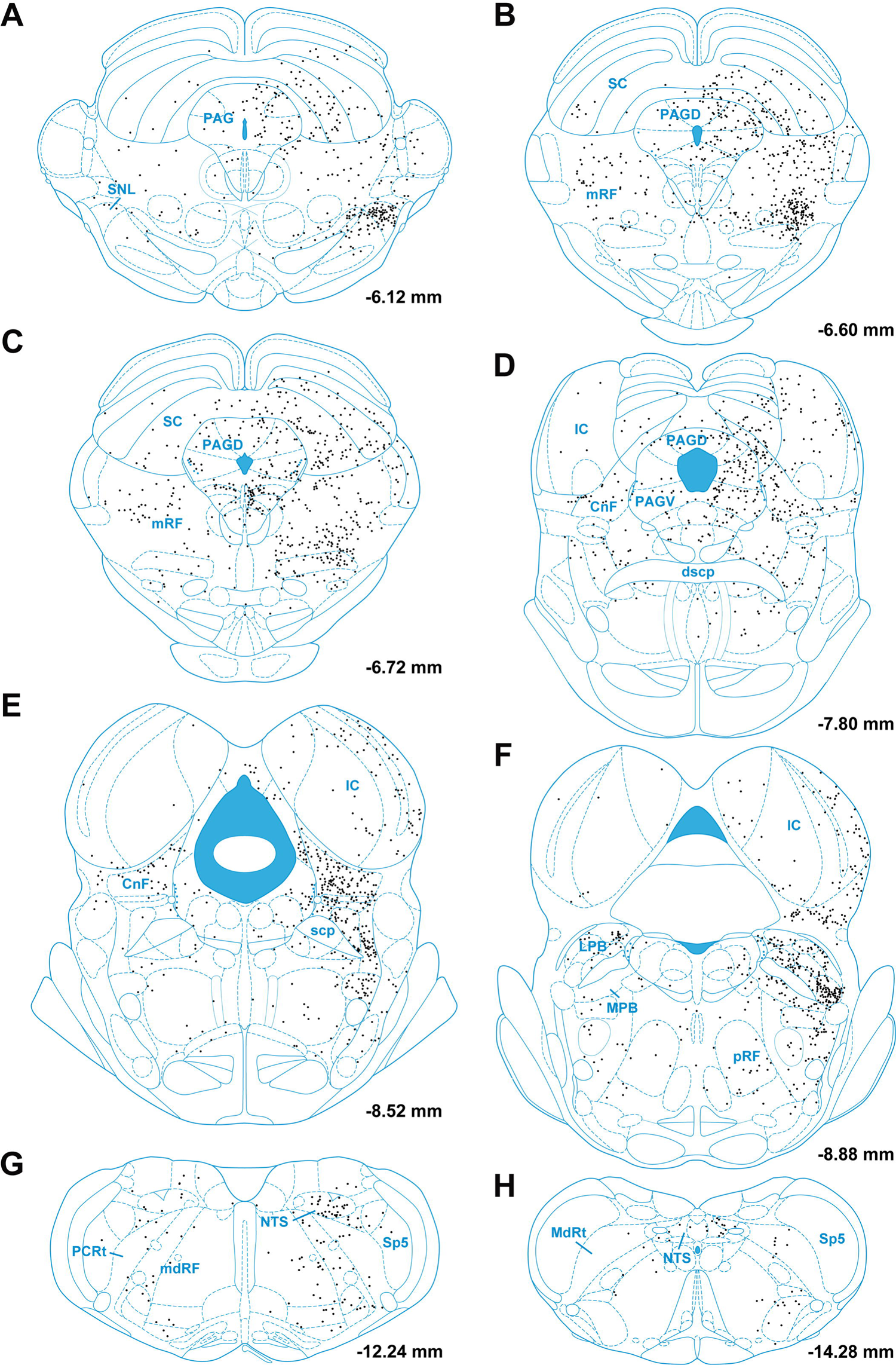
Drawings showing the location of brainstem input cells to LPB-PVT neurons in one representative case (case with starter cells indicated as magenta colored symbols in Fig. 6). The locations of the input cells are shown from anterior to posterior levels of the brainstem (**a-h**). See list for abbreviations and numbers at the bottom represent distance from the bregma.

**Fig 9.**
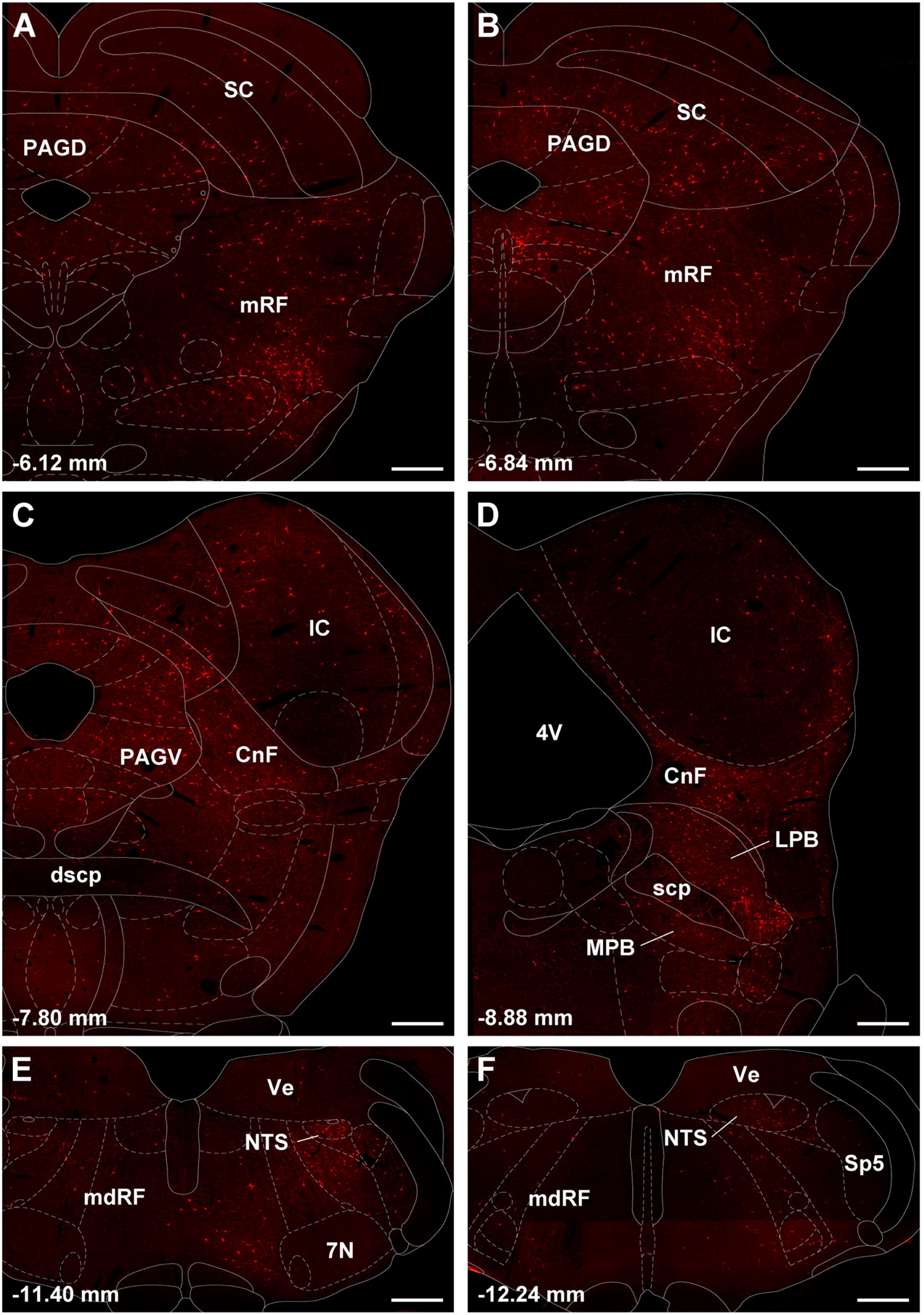
Images of brainstem sections showing mCherry labeling from the TRIO approach applied to LPB-PVT neurons in the case shown in Fig. 8. The sections are shown from the anterior to posterior levels of the brainstem (**a-f**). See list for abbreviations and numbers at the bottom represent distance from the bregma. Scale bars: 500 µm.

The forebrain provided fewer input cells than the brainstem. However, in some instances, the input cells formed tight clusters within regions that were mostly devoid of any other input cells. Especially noteworthy were the clusters observed in the BSTDL (Figs. 10a-b, 11a), CeL (Figs. 10e-f, 11b), and parasubthalamic nucleus (Figs. 10g-h, 11c). Input cells were also found in the paraventricular nucleus of the hypothalamus (Fig. 10d), perifornical region of the lateral hypothalamus (Fig. 10e-f), and the interstitial nucleus of the posterior limb of the anterior commissure (Fig. 10d). Other regions of the forebrain only contain an occasional scattering of input cells and most regions including all of the cortex and striatum were entirely devoid of cells.

**Fig 10.**
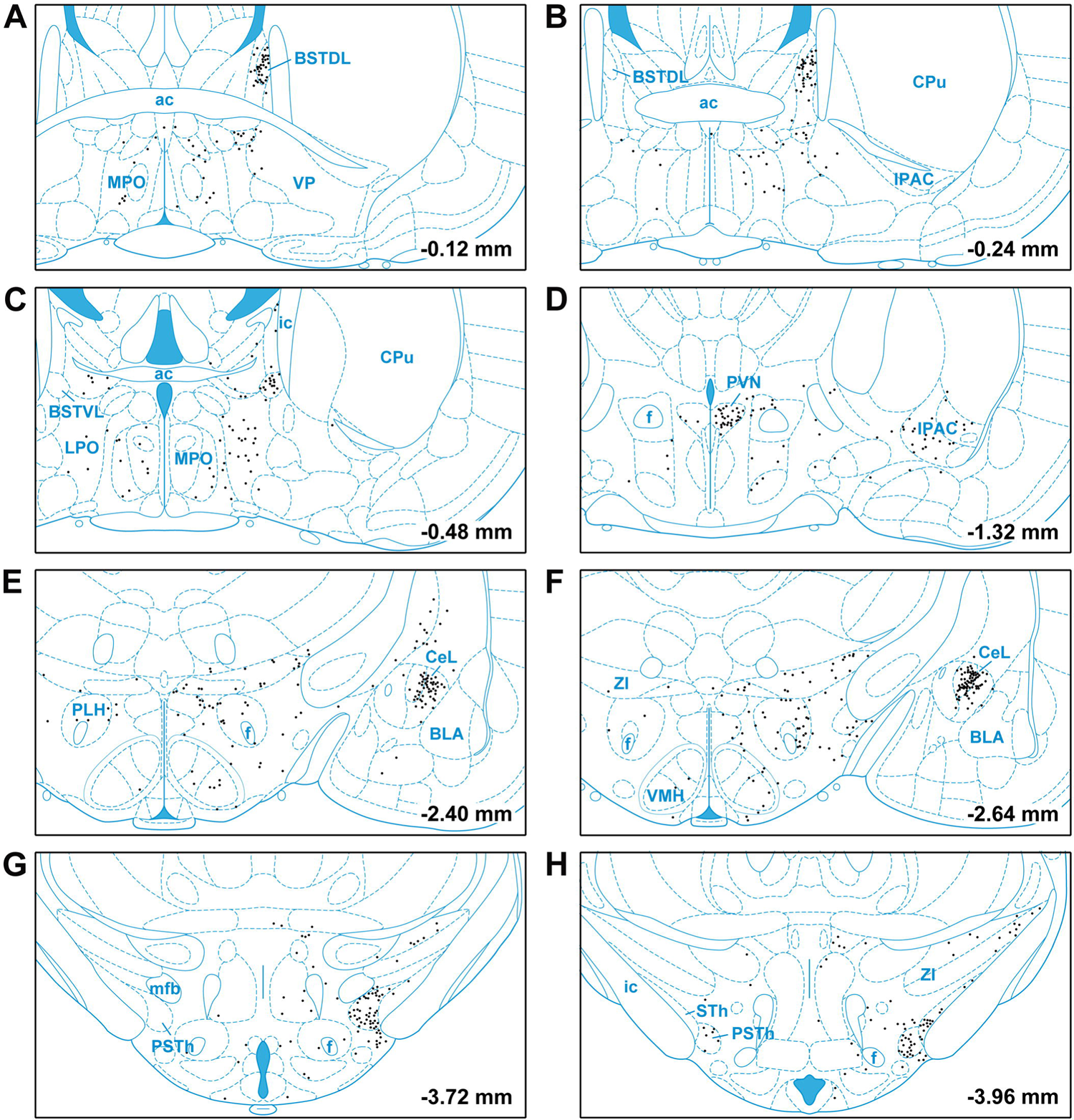
Drawings showing the location of input cells from the forebrain to LPB-PVT neurons. The locations of the input cells are shown from the anterior to posterior most levels of the forebrain (**a-h**). See list for abbreviations and numbers at the bottom represent distance from the bregma.

**Fig 11.**
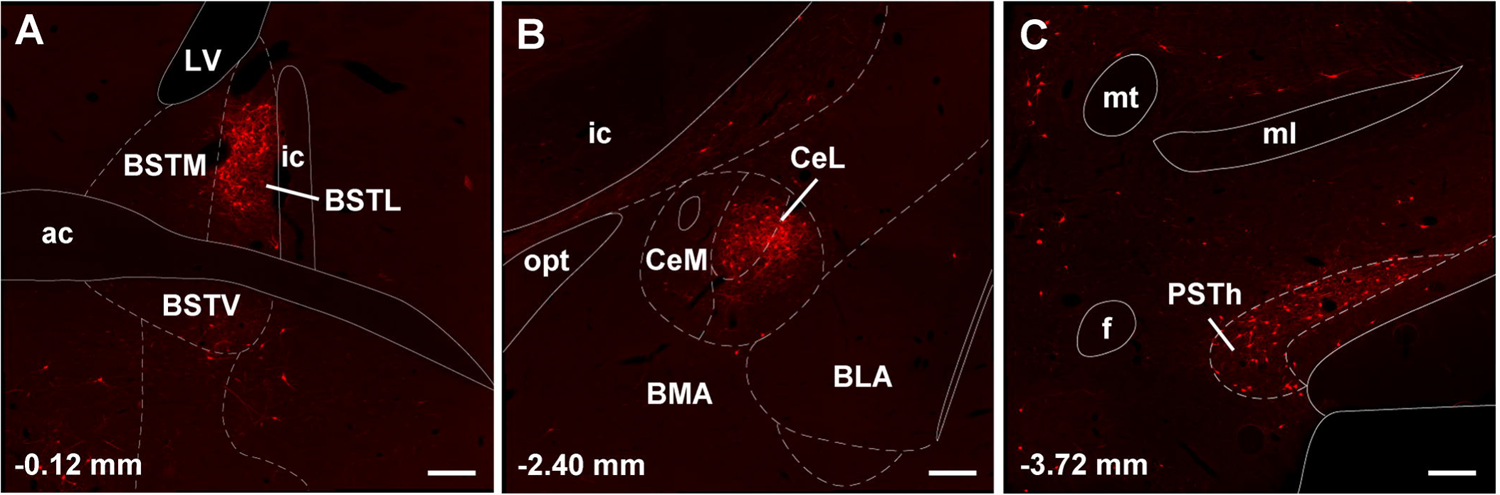
Images of the forebrain showing clusters of mCherry input cells making synapses on LPB-PVT projecting neurons. The sections are from the case shown in Fig. 10 and are arranged from the anterior to posterior most levels of the forebrain (**a-c**). See list for abbreviations and numbers at the bottom represent distance from the bregma. Scale bars: 250 µm.

Starter cells for the experiments involving injections of AAVrg-EBFP-Cre in the CeL (n = 2, 302 and 331 starter cells) were located in the LPB and MPB (Fig. 12a). There were more starter cells in the MPB associated with the CeL than with the PVT experiments. Input cells for PB-CeL experiments (Fig. 12b) were in similar brain regions as for the PB-PVT experiments. One notable exception was a large number of input cells in the reticular portion of the substantia nigra in the PB-CeL cases (Fig. 12c) which was largely absent in the PB-PVT cases (Fig. 8a). Figure 13 shows the proportion of input cells from different regions of the brain associated with the PB-PVT and PB-CeL projecting neurons.

**Fig 12.**
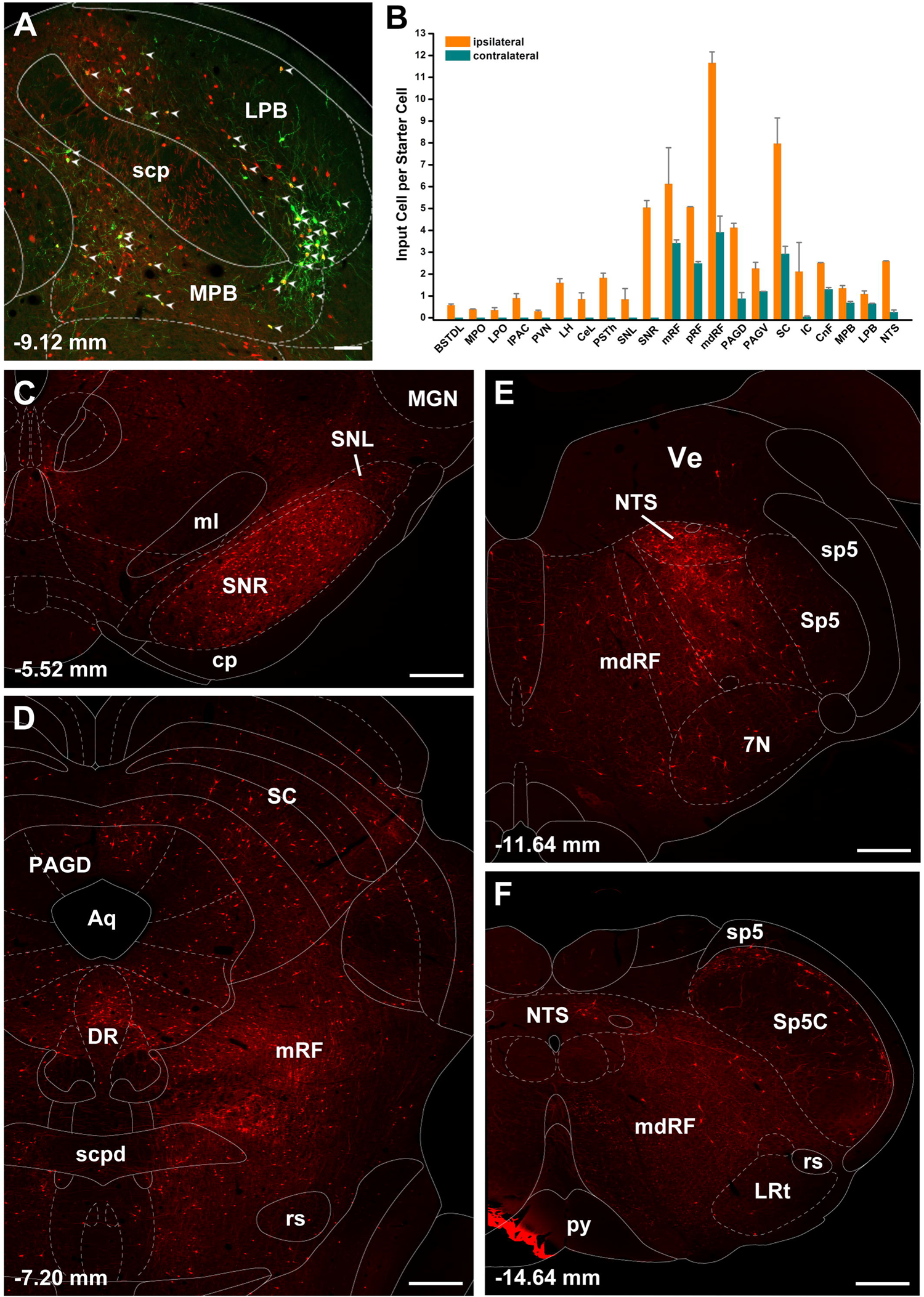
Input cells to the PB-CeL neurons. Image of the PB showing starter cells in one case (**a**) and the number of input cells per starter cells in the brain (**b**). Images of the brainstem showing mCherry input cells to PB-CeL projecting neurons arranged from the anterior to posterior most levels of the forebrain (**c-f**). See list for abbreviations and numbers at the bottom represent distance from the bregma. Scale bars: 100 µm for **a**, 500 µm for **c-f**.

**Fig 13.**
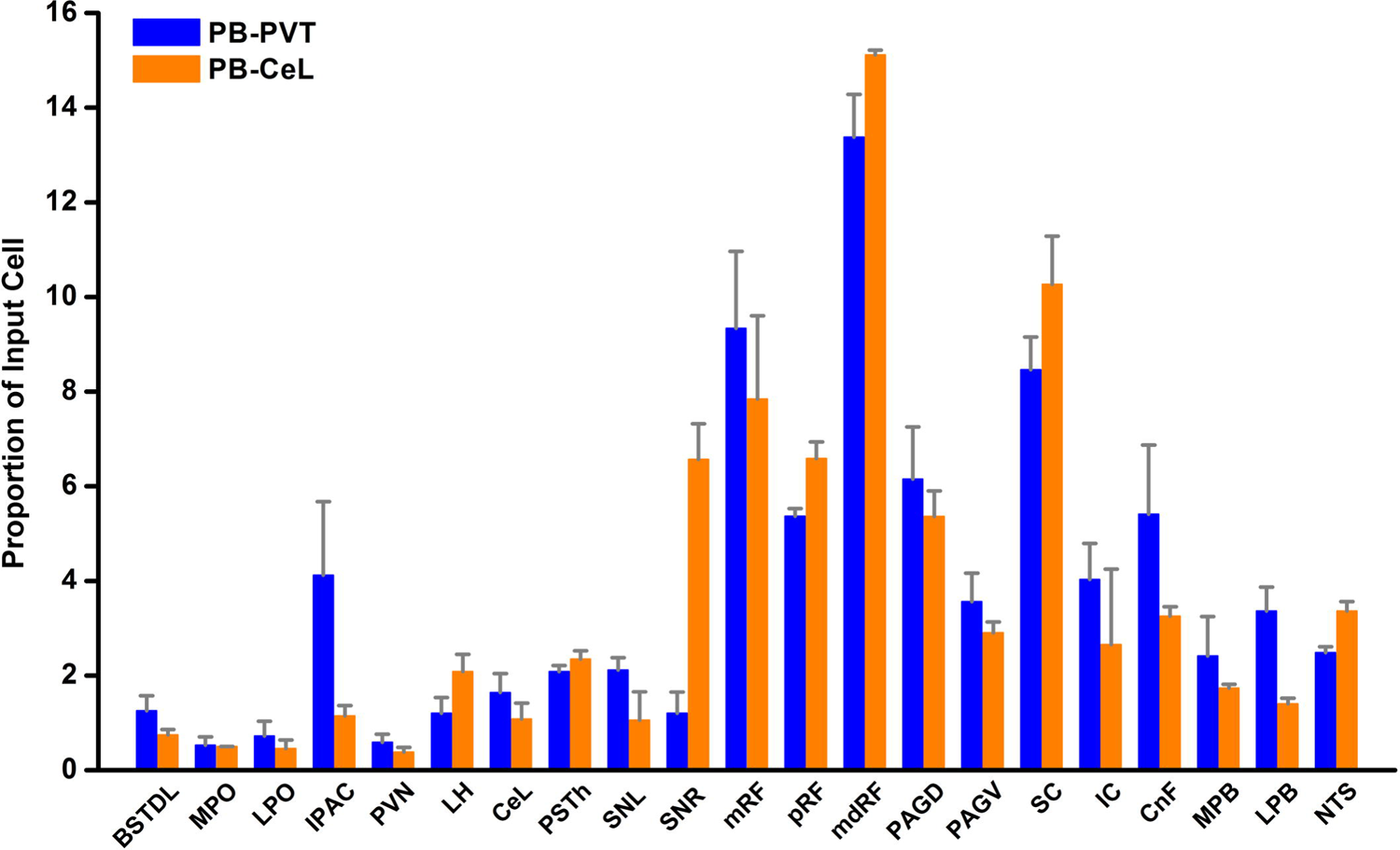
Proportion of input cells to PB neurons that synapse on the PVT and CeL.

Control experiments were done with injections of the helper AAV and RVdG in the PB of rats that had not receive the AAVrg-EBFP-Cre in the PVT since there is always a possibility that RVdG is transduced in neurons that lack Cre (Gehrlach et al. 2020; Miyamichi et al. 2013). We confirm the lack of co-distribution of mCherry and HA labeling in PB neurons and that there was a total absence of mCherry labeling in the entire brain. Additional controls were done with AAVrg-EBFP-Cre in the PVT without helper AAV in the PB followed by RVdG which show the same results as the no-Cre controls. These observations indicate that mCherry cells observed in the cases analyzed are indeed input cells to PB-PVT projecting neurons.

### Intersectional anterograde tracing

Injections of the AAVrg-EBFP-Cre were confined mainly to the aPVT and pPVT as evidenced by the reporter protein EBFP being confined to PVT neurons (Fig. 14a, 14c). These injections along with injections of AAV9/Flex-GFP above the superior cerebellar peduncle at the pontine-midbrain transition resulted in GFP expressing neurons mostly in the LPB in a pattern similar to what was observed in the TRIO tracing experiment (Fig. 14e). Transduced GFP fibers partially overlapped with the expression of EBFP in both the aPVT and pPVT indicating a point to point projection from the LPB to the PVT (Fig. 14b, 14d). A dense plexus of GFP fibers containing enlargements was observed bilaterally in both the aPVT and pPVT (Fig. 14f-g1).

**Fig 14.**
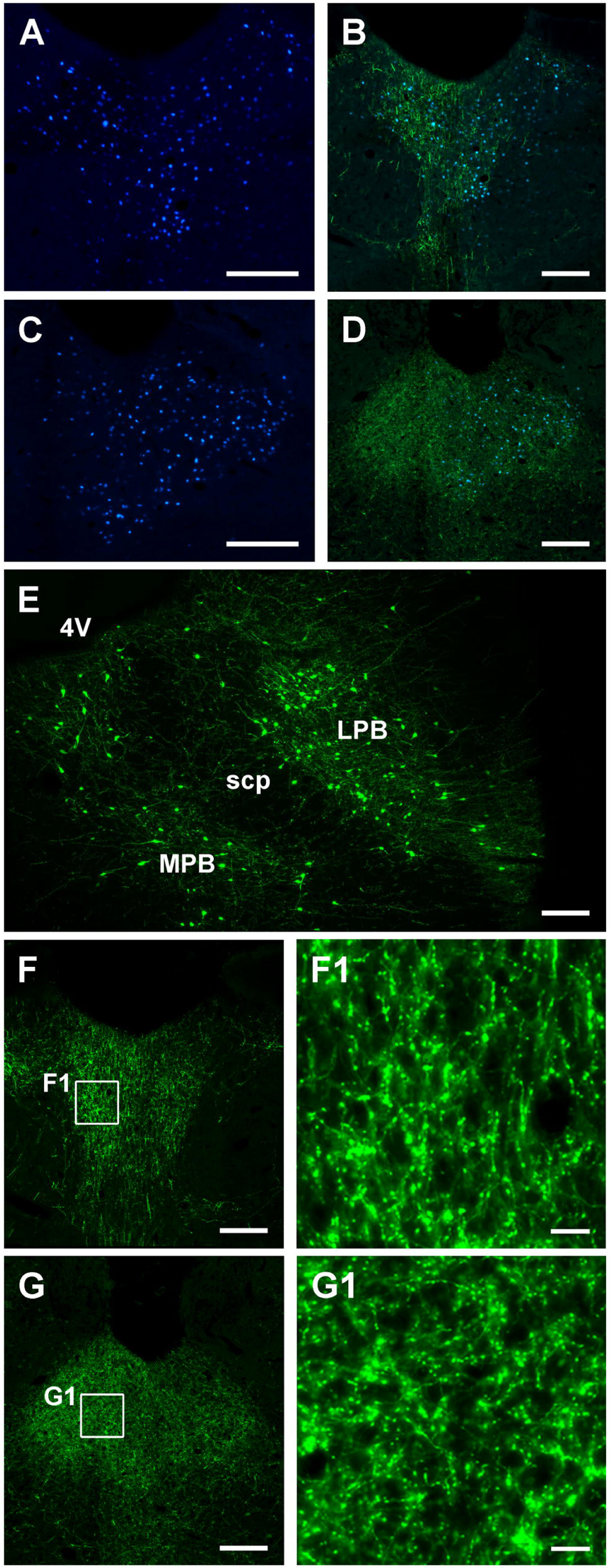
Collateral projections of PB-PVT neurons as shown using the intersectional anterograde tracing approach. Distribution of EBFP (blue) in the aPVT (**a**) and pPVT (**c**) produced by injections of the AAVrg-EBFP-Cre in the PVT with resulting GFP (green) anterograde fiber labeling overlapping the EBFP (blue) for the aPVT (**b**) and pPVT (**d**). Neurons were transduced in most regions of the PB but were concentrated in the LPB (**e**) producing dense GFP fiber labeling in the aPVT (**f**) and pPVT (**g**). Images **f1** and **g1** are higher magnification of the insets in **f** and **g**, respectively. See list for abbreviations and numbers at the bottom represent distance from the bregma. Scale bars: 200 µm for **a-g**, 20 µm for **f1** and **g1**.

Other midline and intralaminar thalamic nuclei contained few GFP fibers (not shown). It is clear from examining the whole brain that PB-PVT neurons have axons that are highly collateralized because a number of areas in addition to the PVT contained a moderately dense plexus of GFP fibers. Fiber labeling was generally more robust on the ipsilateral side although the PVT receive a dense bilateral innervation. Cortical areas including the prefrontal cortex were devoid of fibers whereas the lateral septal area and the diagonal band of broca contained mostly scattered fibers. There was a moderate density of GFP fibers in the median preoptic nucleus (Fig. 15a, 15a1) and anterior aspect of the lateral preoptic area (Fig. 15b, 15b1) whereas the medial preoptic area was sparsely labeled (Fig. 15b). Fibers were seen in the zona incerta where they merged with a moderately dense fiber plexus found in the paraventricular nucleus of the hypothalamus (Fig. 15c, 15c1). The dorsomedial nucleus of the hypothalamus and perifornical lateral hypothalamic area contained a moderately heavy density of fibers (Fig. 15d-e1). Labeled fibers were observed crossing in the retrochiasm with a few of these fibers showing enlargements but with the majority appearing as smooth fibers of passage (arrow, Fig. 15c-e). Some regions of the brain were expected to contain GFP fibers based on previous description of the projections pattern provided by LPB neurons (Huang et al. 2021a). For instance, the BSTDL and CeL were almost completely devoid of fiber labeling except for a few scattered fibers (Fig. 16a, 16b). Ascending fibers of passage were observed in the substantia nigra, ventral tegmental area, retrorubral field, and pontine tegmentum as they extended towards the forebrain. Fibers ascending to the thalamus could also be seen passing along the cerebral aqueduct and the third ventricle whereas fiber labeling in the remainder of the brainstem was sparse except for a few regions. The dorsal and ventral columns of the PAG contained a moderate amount of fibers as did the pontine tegmentum (Fig. 16c). The medulla was largely devoid of labeled fibers except for light labeling in the raphe nuclei above the pyramidal tract (data not shown).

**Fig 15.**
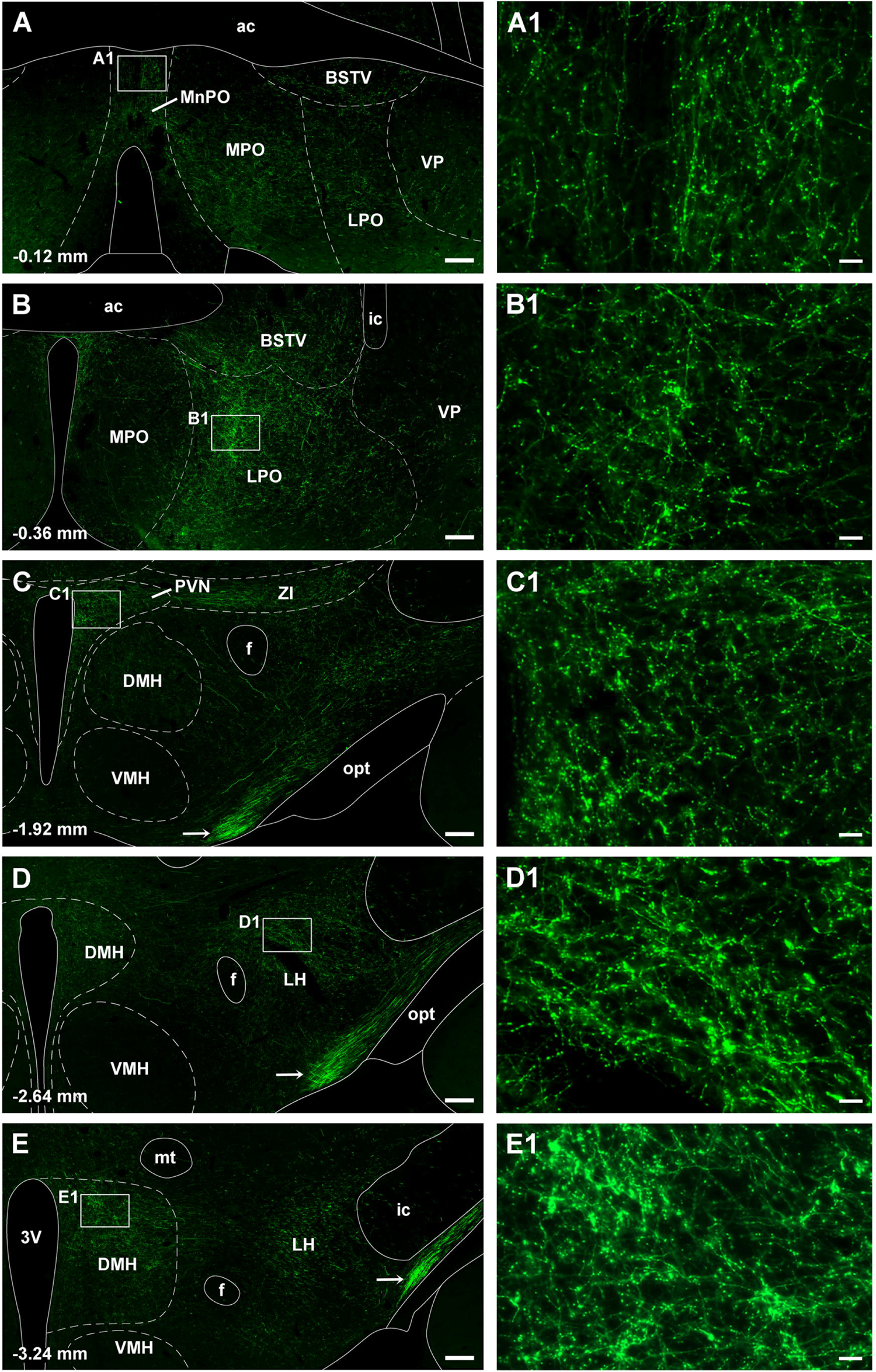
Images of the forebrain containing moderately dense GFP fiber labeling produced by the intersectional anterograde tracing approach applied to the PB-PVT neurons. Images of the GFP labeling produced by the case shown in Fig. 14 from the anterior to posterior levels of the forebrain (**a-e**). Images on the right are higher magnification of the inset shown on the left. See list for abbreviations and numbers at the bottom represent distance from the bregma. Scale bars: 200 µm for **a-e**, 20 µm for **a1-e1**.

**Fig 16.**
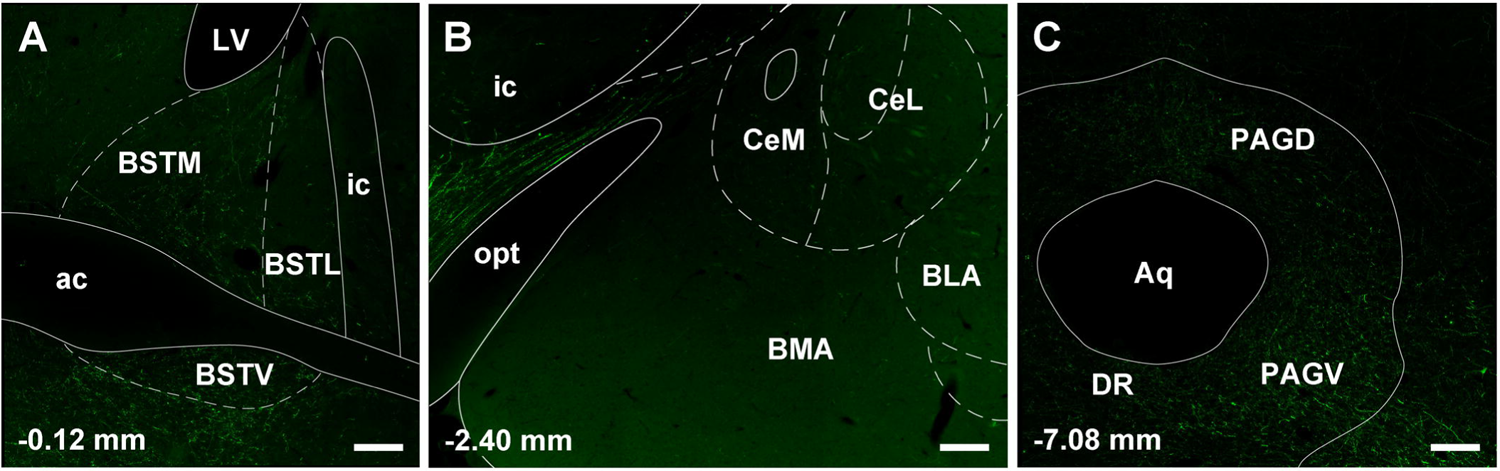
Images of brain regions lacking substantial GFP fiber labeling produced by the intersectional anterograde tracing approach applied to PB-PVT neurons. A relative lack of labeling was observed in the BSTDL (**a**), CeL (**b**) whereas the ventrolateral region of the PAG contained weak fiber labeling (**c**). See list for abbreviations and numbers at the bottom represent distance from the bregma. Scale bars: 200 µm.

## Discussion

The results of the CTB tracing experiments are in line with previous studies reporting that a relatively small number of neurons associated with catecholamine cell groups project to the PVT (Cornwall and Phillipson 1988; Chen and Su 1990; Krout et al. 2002; Krout and Loewy 2000a, b; Krout et al. 2003; Li and Kirouac 2012; Phillipson and Bohn 1994; Otake et al. 1994). The relatively large number of neurons in the LPB compared to the rest of the brainstem is also consistent with our quantitative analysis of retrograde labeling in the brain following small restricted injections of CTB in the aPVT and pPVT (Li and Kirouac 2012). It is generally understood that the PB relays brainstem and spinal cord viscerosensory, somatosensory and taste related signals to widespread regions of the forebrain (Chiang et al. 2019; Palmiter 2018; Herbert et al. 1990; Saper and Loewy 1980) including the PVT (Li and Kirouac 2012; Krout and Loewy 2000a). Mapping of brain-wide sources of neurons that make synapses on PB-PVT projecting neurons in the TRIO experiments revealed that the largest source of input cells is from neurons that are scattered in the brainstem reticular formation. Other prominent sources of input cells include the PAG in addition to the superior and inferior colliculi of the midbrain tectum. Another notable observation from the rabies TRIO experiments is that the BSTDL and CeL are distinctive sources of afferents to PB-PVT neurons. The intersectional anterograde tracing approach demonstrates that PB neurons that project to the PVT do not innervate the BSTDL and CeL but provide collateral innervation of a number of forebrain regions postulated to regulate wakefulness and homeostasis. As discussed below, these anatomical findings provide evidence that the PVT is part of an interconnected network of neurons dispersed in the brainstem and forebrain that is involved in the regulation of arousal and behavioral states.

### Lower brainstem inputs to the PVT

The PVT contains a relatively sparse plexus of TH fibers (Takada et al. 1990; Otake and Ruggiero 1995) which has been shown to mostly originate from dopamine neurons in the hypothalamus and PAG (Li et al. 2014). The number and location of TH neurons in the lower brainstem that project to the aPVT and pPVT were assessed to further characterize the source of these fibers. A relatively small number of TH neurons confined to the C2/A2 cell group of the NTS and the C1/A1 cell group of the caudal VLM were found to project to the PVT as previously reported (Phillipson and Bohn 1994; Otake et al. 1994). Neurons that stain for TH may synthesize either norepinephrine or epinephrine depending on the presence of other rate-limiting enzymes (the nomenclature and function of the catecholamine cell groups are discussed in Bucci et al. 2017). Consistent with our previous retrograde tracing experiments (Li and Kirouac 2012; Li et al. 2014; Kirouac et al. 2006), there was a paucity of CTB cells in the LC even though a few projection neurons has been reported by others (Krout et al. 2002; Otake and Ruggiero 1995).

One likely explanation for this discrepancy may be that our approach of using fine glass pipettes to administer a small volume of CTB limited the amount of the tracer that was picked up by fibers coursing through the area. Viral transduction of LC neurons in dopamine β Cre mice provided evidence that the LC produces and releases dopamine in the PVT (Beas et al. 2018), but a functional LC projection to the PVT is difficult to envision considering the results of our retrograde tracing experiments and the view that neuronal projections are well conserved between rats and mice (Ellenbroek and Youn 2016).

The brainstem catecholamine cell groups and the nuclei with which they are associated are implicated in the viscerosensory reflexes and the execution of various behavioral and physiological responses to homeostatic challenges (Guyenet 2006; Guyenet et al. 2013; Ritter et al. 2006; Rinaman 2011). Consequently, it is possible that catecholamine signals may be integrated by the PVT to promote adaptive responses, as recently shown for hypoglycemia-induced food intake by a catecholamine VLM-PVT projection (Sofia Beas et al. 2020). However, the type of influence catecholamines have on PVT functions remains an open question for a number of reasons. First, catecholamine neurons that innervate the PVT are widely scattered in both the hypothalamus and brainstem and do not appear to form functionally distinct populations. Second, experimental evidence indicates that catecholamine neurons in the lower brainstem exert cellular effects predominately via conventional glutamatergic transmission (DePuy et al. 2013) and potentially the co-release of neuropeptides but not catecholamines (Guyenet et al. 2013).

Third, the fact that brainstem catecholamine neurons form a meshwork of interconnected neurons that provide highly collateralized axons indicates that these neurons are likely to produce broad and diffused arousal effects on the brain including effect via extrasynaptic release and volume transmission (Aston-Jones and Cohen 2005; Bucci et al. 2017; Card et al. 2006; Guyenet et al. 2013; Patthy et al. 2021; Fuxe et al. 2010).

The PB is a multisensory relay center that has been subdivided into medial and lateral regions that have been further partitioned into cytoarchitecturally distinct subnulei (Chiang et al. 2019; Saper and Loewy 1980). It is clear from the present and our previous retrograde tracing experiments that the LPB is the area of the brainstem that contains the largest number of PVT projecting neurons (for quantitative analysis of the source of all brain inputs to the PVT, see Li and Kirouac 2012). We did not attempt to localize PB-PVT projecting neurons within subnuclear boundaries because of the difficulty of doing this with precision in thick sections stained with NeuN. The fact that the dendrites of LPB neurons extend outside subnuclear boundaries further makes the functional relevance of these anatomical demarcations debatable (Sarhan et al. 2005; Jasmin et al. 1997). However, it is clear that CTB neurons were located in the general region of the LPB that contains the external, ventral and dorsal lateral subnuclei that receive afferents from general viscerosensory integration regions of the NTS (Chiang et al. 2019; Saper and Loewy 1980; Herbert et al. 1990) and nociceptive signals from the brainstem and spinal cord (Chiang et al. 2019; Jaramillo et al. 2021; Han et al. 2015; Gauriau and Bernard 2002; Deng et al. 2020). It is also apparent that most of the PVT projecting neurons are localized in the LPB, and for this reason, we will refer to them as LPB-PVT neurons with the caveat that some neurons in the MPB contribute to this projection.

### Whole brain inputs to LPB-PVT projecting neurons

It was expected that the NTS would be the major source of input cells to LPB-PVT neurons because of the existence of a dense projection from the NTS to the LPB (Herbert et al. 1990; Moga et al. 1990; Tokita et al. 2009). Nonetheless, we observed a low number of cells in the NTS compared to the large number input cells scattered broadly throughout the brainstem RF. This highlights the clear advantage of the rabies monosynaptic approach over traditional tracing methods that simply assess area-to-area projection patterns and density but not if the neurons are synaptically connected. This is especially important for LPB neurons which have dendrites that extend hundreds of micrometers (Sarhan et al. 2005). It should be appreciated that the large number of input cells in the RF represent cells that are distributed over a large span of neural tissue and not as smaller clusters of cells like observed in the BSTDL, CeL and NTS. The brainstem RF was initially conceptualized as a loosely organized network of interconnected neurons that integrate polymodal inputs and coordinate many vital functions (Blessing 1997). In addition, the brainstem RF is also postulated to be part of the ascending reticular activating system involved in the maintenance of arousal and behavioral states (Blessing 1997; Moruzzi and Magoun 1949; Liu and Dan 2019; Jones 2020; Scammell et al. 2017). Some regions of the RF are regarded as nuclei with sensory, motor or mixed functions (Blessing 1997; Jones 1995).

Except for a few noteworthy examples, clear and distinct clusters of input cells were generally not found in most of the anatomically recognized nuclei in the brainstem. First, the NTS contained a relatively small number of input cells located primarily anterior to the beginning of the fourth ventricle. The neuroanatomy of the NTS is complex and it is impossible to know the exact types of signals that these input cells transmit simply based on their location in the NTS (Herbert et al. 1990; Lundry and Norgren 2015; Blessing 1997). The fact that the anterior portions of the NTS receive gustatory afferents from the tongue in addition to overlapping viscero- and somato-sensory afferents from the oral cavity (Herbert et al. 1990; Lundry and Norgren 2015; Blessing 1997) suggests that LPB-PVT neurons may receive integrated polymodal sensory information from the oral cavity via the NTS. However, taste is not likely the predominant signal received by LPB-PVT because these neurons are localized primarily in the nongustatory regions of the PB (Blessing 1997; Chiang et al. 2019; Herbert et al. 1990). Second, the parvicellular reticular nucleus of the medulla located adjacent to the anterior most portion of the NTS contained a distinctive collection of input cells. The parvicellular reticular nucleus processes signals related to sensory and motor functions associated with oral movements (Travers et al. 2000; Zerari-Mailly et al. 2001; Moriyama 1987; Takatori et al. 1981; Shammah-Lagnado et al. 1992) pointing to the possibility that the LPB-PVT neurons may receive signals related to ingestion or emesis. The absence of input cells in the nociceptive superficial lamina of the spinal trigeminal nucleus suggests that LPB-PVT neurons may not directly receive nociceptive signals from the head. We cannot exclude the possibility that LPB-PVT neurons receive nociceptive signals from the spinal cord since sections beyond the caudal medulla were not examined.

Input cells were scattered across multiple stereotaxic levels in a number of midbrain regions. These included the PAG, involved in the regulation of emotional and nociceptive responses (George et al. 2019; Behbehani 1995); the superior and inferior colliculi, implicated in the modulation of movements and arousal to visual and auditory stimuli (Brandao et al. 1994; Brandao et al. 2003; Schenberg et al. 2005; Isa et al. 2021; Mulckhuyse 2018; Basso and May 2017; Cabrera et al. 2013); and the nucleus cuneiformis, linked to the modulation of locomotion and nociception (Coles et al. 1989; Jordan et al. 2008; Ebrahimzadeh and Haghparast 2011; Haghparast et al. 2010; Sandner et al. 1993Bernard, 1989 #3859). The input cells were not concentrated within distinctive subregions of the PAG or tectum suggesting that LPB-PVT neurons may be integrating nonspecific signals from these regions.

Several forebrain areas linked to the regulation of homeostasis and modulation of behavioral states contained a small number of input cells. First, input cells were found scattered in the median preoptic nucleus and the lateral preoptic area where some neurons display enhance activity during sleep (McKinley et al. 2015; Szymusiak et al. 2007; Scammell et al. 2017). Sleep actives neurons in the preoptic area contain GABA (Gong et al. 2004) which are believed to promote sleep states by inhibiting arousal centers in the hypothalamus and brainstem (Szymusiak et al. 2007; Scammell et al. 2017). In addition to the regulation of sleep, the median preoptic nucleus is connected with areas of the hypothalamus involved in the regulation of body fluids, sodium, and temperature (McKinley et al. 2015; Menani et al. 2014). Input cells were also consistently observed in the LH, a functionally complex region that is involved in behavioral and physiological responses associated with survival including the regulation of sleep, arousal, energy balance, reward, stress and defensive responses (Berthoud 2002; Castro et al. 2015; Arrigoni et al. 2019; Scammell et al. 2017; Alexandre et al. 2013; Bonnavion et al. 2016).

Input cells were found as distinct clusters in the BSTDL, CeL, IPAC and parasubthalamic nucleus. The BSTDL, CeL and IPAC form a macrostructure called the extended amygdala with common anatomical and functional characteristics (de Olmos et al. 2004; Alheid 2003; Alheid et al. 1999). The parasubthalamic nucleus is interconnected with the BSTDL, CeL, IPAC and LPB; and is postulated to be involved in the execution of integrated behavioral and physiological responses (Goto and Swanson 2004; Shah et al. 2021), as recently demonstrated for fear-induced hypothermia (Liu et al. 2021a). The parasubthalamic nucleus also projects to the pPVT (Goto and Swanson 2004) and regions of the prefrontal cortex that in turn innervate the pPVT and the extended amygdala (Li and Kirouac 2012). The BSTDL/CeL are best known for their role in regulating responses to threats (Lebow and Chen 2016; Gungor and Pare 2016; Duvarci and Pare 2014; Fox et al. 2015; Davis and Shi 1999), potentially in response to feedforward signals about aversive conditions from the LPB (Han et al. 2015; Phua et al. 2021; Cai et al. 2018; Jaramillo et al. 2020; Ito et al. 2021; Bowen et al. 2020; Chiang et al. 2020) and the PVT (Do-Monte et al. 2017; Do-Monte et al. 2015; Penzo et al. 2015; Pliota et al. 2018). Subpopulations of GABA neurons in the BSTDL/CeL express unique neurochemical markers that differentially contribute to defensive responses through mechanisms and projections that are not completely understood (Gungor and Pare 2016; Duvarci and Pare 2014; Fadok et al. 2018; Li 2019). Regardless of neuropeptides associated with these neurons, they are likely to have inhibitory effects on LPB-PVT neurons because GABA is the primary neurotransmitter associated with these projection neurons (Moga and Gray 1985; Ye and Veinante 2019:Bartonjo, 2020 #3833). Although we did not identify the neuropeptide associated with the neurons in the BSTDL/CeL that synapse on LPB-PVT neurons, corticotropin-releasing factor, somatostatin, and neurotensin are potential candidates as these markers have been shown to be associated with some BSTDL/CeL GABAergic neurons that project to the LPB (Moga and Gray 1985; Ye and Veinante 2019:Bartonjo, 2020 #3833). Future investigation on the identity and activation conditions of BSTDL/CeL neurons that synapse on LPB-PVT neurons would yield critical information about how these long-range circuits function in the brain.

The lateral substantia nigra (SNL) is another brain region found to provide substantial input to LPB-PVT neurons. This group of cells is of special interest because it receives afferent input from the BSTDL/CeL (Steinberg et al. 2020; Gonzales and Chesselet 1990; Vankova et al. 1992; Liu et al. 2021b) and a projection from the CeL to the SNL has been shown to modulate both appetitive and aversive learning (Steinberg et al. 2020). Furthermore, the SNL and LPB are reciprocally connected (Tokita et al. 2009; Huang et al. 2021a; Vankova et al. 1992) forming potentially an interconnected network of neurons involving the PB, PVT, SNL and BSTDL/CeL. Likewise, the LPB may also indirectly send neural signals to the BSTDL/CeL via projections to the PVT and parasubthalamic nucleus (Li and Kirouac 2012; Huang et al. 2021a; Huang et al. 2021b) which in turn densely innervate the BSTDL/CeL (Li and Kirouac 2008; Goto and Swanson 2004) forming another interconnected network involving the LPB, PVT and the BSTDL/CeL.

The location and density of input cells observed in the TRIO experiments were generally consistent with what has been reported in rats and mice following injections of retrograde tracers involving much of the PB (Moga et al. 1990; Tokita et al. 2009; Herbert et al. 1990). However, there were notable exceptions including the absence of input cells from prefrontal cortical areas and medial region of the central nucleus of the amygdala. In fact, the large number of input cells in the CeL combined with the absence of such cells in the medial part of the central nucleus attests to the specificity of the synaptic connections associated with a CeL-PB-PVT circuit. It is clear that the synaptic inputs to LPB neurons from the BSTDL/CeL are not unique to neurons that project to the PVT since the TRIO approach applied to PB neurons that project to the CeL also resulted in distinctive clusters of input cells in the BSTDL/CeL. The sources of inputs to PB-PVT and PB-CeL neurons were similar except for the reticular portion of the substantia nigra that appeared to preferentially synapse on PB-CeL neurons. This region of the substantia nigra is a basal ganglia output nucleus involved in the regulation of motor and cognitive responses via thalamocortical feedback circuits (Shipp 2017; Alexander et al. 1990; Deniau et al. 2007).

### Whole brain projections of LPB-PVT neurons

Anterograde tracing of the collateral innervation pattern of LPB-PVT neurons indicates that these neurons are largely specific to the PVT although retrograde tracing studies indicate that many midline thalamic nuclei receive input from the LPB (Krout et al. 2002; Krout and Loewy 2000a). The present paper and other recent evidence support the view that some neurons in the PB preferentially innervate unique thalamic nuclei (Deng et al. 2020; Huang et al. 2021a).

Previous retrograde tracing studies have shown that some LPB neurons that express unique peptides, including CRF, substance P, cholecystokinin (CCK) and cocaine- and amphetamine-regulated transcript (CART), project to the PVT (Kirouac et al. 2006; Otake and Nakamura 1995; Otake 2005). Experimental approaches targeting marker-specific neurons have been used to identify subpopulations of function- and projection-specific neurons in the PB. There are a number of examples in which the evidence indicates that subpopulations of LPB neurons that express particular peptides may be differentially activated by particular homeostatic challenges including nociception, body temperature, taste stimuli, hypoxia and other life threatening states (see reviews by Jaramillo et al. 2021; Chiang et al. 2019; Palmiter 2018). For example, neurons that express calcitonin gene-related peptide (CGRP) have been shown to be activated by most threats examined (Carter et al. 2013; Han et al. 2015; Palmiter 2018) and to project robustly to the BSTDL and CeL (Huang et al. 2021b). However, it is apparent from our anterograde tracing experiments that LPB neurons that innervate the PVT are different from the population that projects to the BSTDL/CeL. Neurons that express dynorphin have been shown to respond to changes in ambient temperatures and to have projection patterns different from the CGRP cells (Geerling et al. 2016; Huang et al. 2021a). Other research groups have shown that dynorphin neurons in the LPB respond to a variety of somatosensory signals including nociception relayed from local CGRP neurons (Choi et al. 2020; Kim et al. 2020; Luskin et al. 2021; Chiang et al. 2020). Glucose-sensing PB neurons that express CCK were shown to project to the ventromedial nucleus of the hypothalamus but not the PVT (Garfield et al. 2014). No attempt was made in the present study to identify a unique neuropeptide associated with LPB-PVT neurons because of the complexity in the number of neurons in the PB that express one or more peptide (Palmiter 2018; Chiang et al. 2019; Zhu et al. 2022). Regardless if LPB-PVT neurons have an unique neuropeptide identity, PB neurons use glutamate as a primary neurotransmitter exert excitatory effects on PVT neurons (Huang et al. 2021a; Geerling et al. 2017; Niu et al. 2010 Zhu, 2022 #3955). Nevertheless, application of single-cell sequencing to categorize unique genes highly expressed in LPB-PVT neurons will be helpful for examining the function of these neurons.

The collateral projections provided by LPB-PVT neurons hint at potential functions associated with these neurons. Areas receiving most significant collateral innervation included the preoptic area, zona incerta, LH, and dorsomedial nucleus of the hypothalamus (DMH). Many of these regions are associated with arousal and the regulation of behavioral states. The zona incerta is postulated to be part of a larger brain-wide interconnected network involved in modulation of arousal, attention, and movements including approach and avoidance to interoceptive and exteroceptive sensory information (Mitrofanis 2005). It is notable that the zona incerta is interconnected with many of the same midbrain regions (Kolmac et al. 1998) shown in the TRIO experiments to make synapses on LPB-PVT neurons. The paraventricular nucleus of the hypothalamus is a key integrative and motor nucleus involved in the regulation of the autonomic nervous system and the stress response (Dampney 1994; Myers et al. 2014). The DMH is linked to the modulation of circadian and biological rhythms in addition to contributing to the physiological and hormonal responses to threats (DiMicco et al. 2002; Fontes et al. 2011; Saper et al. 2005; Myers et al. 2014). Finally, and as discussed above, the preoptic area and LH are brain regions that are broadly associated with arousal and the modulation of homeostasis and behavioral states. It is of potential significance that some forebrain regions that contain neurons that make monosynaptic connections with LPB-PVT neurons are the same regions that received significant collateral fiber innervation from LPB-PVT neurons (e.g., LH). While we demonstrate that neurons in the preoptic area and LH make synaptic connections with LPB-PVT neurons, we do not know if fibers from LPB-PVT neurons make synaptic connections with neurons in these reciprocally innervated regions. It is also remarkable that the median preoptic nucleus, DMH, and LH are well-characterized sources of afferents to the PVT (Kirouac et al. 2005; Li and Kirouac 2012; Thompson et al. 1996; Goto and Swanson 2004), further highlighting the possibility that neurons in the PVT may be part of a larger network of neurons with similar functions.

### Functional considerations

This paper presents anatomical evidence that demonstrates that the LPB is the major source of lower brainstem afferents to the PVT. The LPB has received considerable interest for its role in regulating behavioral and physiological responses to exteroeceptive and interoceptive signal related to homeostatic challenges (Palmiter 2018; Jaramillo et al. 2021; Chiang et al. 2019). The fact that the LPB is considered a major integrating and relay center of spinal cord and brainstem sensory signals to forebrain has provided the drive for identification of projection specific neurons in the LPB that regulate behavioral responses to specific challenges. For instance, arousal from sleep during hypercapnia (high CO_2_) was shown to be mediated by CGRP neurons in the LPB that act on circuits primarily located in the basal forebrain and to a lesser extent in the LH and CeL (Kaur et al. 2017). Furthermore, neurons in the LPB that express CGRP also respond to nociception associated with footshocks and conditions known to suppress appetite; and optogenetic activations of these projections to the CeL diminish food intake (Carter et al. 2013) and produce an aversive teaching signal that results in a retrievable fear memory (Han et al. 2015). Neurons in the LPB send highly collateralized fiber projections to interconnected sites in the brain (Sarhan et al. 2005; Bowen et al. 2020; Chiang et al. 2020; Zhu et al. 2022) and attempts at isolating specific defensive behavioral and physiological responses to one anatomical terminal field have proven difficult using optogenetic approaches (Bowen et al. 2020; Chiang et al. 2020). The difficulty in assigning specific functions to subpopulations of PB neurons is compounded by the fact that these neurons exist as intermixed populations and not always as distinct clusters of cells (Huang et al. 2021a; Huang et al. 2021b; Bowen et al. 2020; Barik et al. 2018; Chiang et al. 2020). The latter is further complicated by the fact that neurons in the PB have dendrites and axons that extend across subnuclear zones (Sarhan et al. 2005; Jasmin et al. 1997; Chiang et al. 2020). Another notable observation made in the present paper is that the LPB neurons that innervate the PVT do not project to the BSTDL/CeL. Indeed, our understanding of how the PB integrates and relays signals to other regions of the brain is preliminary and limited by the complexity of the types of neurons that make up the PB including their unique afferent and efferent connections.

The number and location of input cells that synapse on LPB-PVT neurons provide some indications of the type of information these neurons broadcast to the PVT. The fact that major sources of input originate from neurons scattered in brainstem regions functionally associated with general arousal suggests that LPB-PVT projecting neurons may be part of an ascending arousal system. The inputs from the superior and inferior colliculi also indicate that LBP-PVT neurons may integrate signals linked to arousing visual and auditory stimuli. It is generally appreciated that arousal and the maintenance of wakefulness are mediated by a highly distributed network of interconnected neurons that span the forebrain, midbrain, and hindbrain (Liu and Dan 2019; Jones 2020; Scammell et al. 2017). The PB has been identified as a critical brainstem structure involved in arousal from sleep via connections to the basal forebrain and the LH (Venner et al. 2019; Qiu et al. 2016; Fuller et al. 2011; Xu et al. 2021; Kaur et al. 2013; Kaur et al. 2017). While there is some evidence that the PVT is involved in arousal from sleep (Hua et al. 2018; Matyas et al. 2018; Ren et al. 2018), other evidence indicates that the thalamus is not critical for the PB’s cortical arousal effects (Qiu et al. 2016; Fuller et al. 2011. Xu, 2021 #3838).

General arousal has also been conceptualized as a brain activation phenomena that invigorates behavior by enhancing the activity of the sensory and motor circuits (Pfaff et al. 2008; Pfaff 2006). Accordingly, another possibility is that general arousal signals enhance the activity of PVT neurons thereby invigorating behavioral responses. It should also be appreciated that factors that promote behavioral responses have general arousal effects on the brain (Pfaff 2006). Determinants of behavior such as physiologic states (e.g., hunger, thirst, nociception, pheromones) and cues that signal potential appetitive and aversive outcomes are often reported to increase the activity of PVT neurons (for details and references, seeBarson et al. 2020; Millan et al. 2017; Hsu et al. 2014; Kirouac 2015, 2021; Penzo and Gao 2021). We postulate that PVT neurons integrate incoming signals that sense the arousal state of the brain with neural inputs signaling the presence of physiologic states and emotionally salient cues. There are a number of recent observations that support this hypothetical view of PVT function. For instance, the PVT uses hypothalamic and brainstem signals related to the energy and hydration needs to modulate behavioral responses to maintain homeostasis (Livneh et al. 2017; Zhang and van den Pol 2017; Hua et al. 2018; Sofia Beas et al. 2020; Leib et al. 2017; Labouebe et al. 2016; Zhu et al. 2018; Meffre et al. 2019). The PVT also receives signals related to predator odor via the ventromedial nucleus of the hypothalamus to suppress food reward seeking (Engelke et al. 2021). Other evidence shows that the PVT integrates signals related to emotionally salient cues from the prefrontal cortex to modulate behavioral responses (Otis et al. 2019; Otis et al. 2017; Do-Monte et al. 2015; Penzo et al. 2015). The type of influence the PVT has appears to be related to whether a response poses a threat or has a chance of being unproductive (i.e., motivational conflict) supporting the view that the PVT is also involved in the selection of responses based on the integration of a wide range of afferent inputs (Cheng et al. 2018; Do-Monte et al. 2017; Lafferty et al. 2020; Choi et al. 2019; Choi and McNally 2017). As a whole, various experimental and anatomical observations suggest that the PVT integrates a broad range of signals related to saliency of cues, homeostatic state, and general arousal to modulate behavioral responses that promote survival.

Figure 17 summarizes some of the more salient anatomical findings reported in this paper. As discussed above, the number of neurons scattered in the brainstem RF that provide synaptic input to the LPB-PVT neurons is especially noteworthy. In addition, a considerable proportion of inputs originate from neurons located in the PAG and the tectum. Another intriguing source of inputs originates from clusters of neurons in the BSTDL and CeL, areas of the extended amygdala also densely innervated by the PVT (Dong et al. 2017; Li and Kirouac 2008) and known to modulate behavioral responses to threats (Gungor and Pare 2016; Duvarci and Pare 2014; Fadok et al. 2018; Li 2019). The BSTDL/CeL could be part of a feedback circuit that influences the transmission of a variety of signals from the LPB to the PVT (i.e., gating mechanism). It is not known if PVT neurons make synaptic contact with neurons in the BSTDL/CeL that synapse on the LPB-PVT neurons nor do we know if the PVT neurons that project to the BSTDL/CeL receive synaptic contacts from the LPB neurons. Nonetheless, this is an intriguing possibility as it would provide a means by which a multisynaptic circuit could modulated the flow of arousal and/or other homeostasis signals to the PVT and other areas of the forebrain. In support, a recent paper identified projections from the BSTDL that modulate threat assessment and feeding by acting on PB neurons (Luskin et al. 2021). Both the LPB and PVT are areas of the brain often reported to be activated during aroused and emotional states (Hsu et al. 2014; Kirouac 2015; Palmiter 2018; Jaramillo et al. 2021; Chiang et al. 2019). Optogenetic excitation of PB fibers in the PVT and chemogenetic activation of PB-PVT neurons elicit responses indicative of an aversive state (Zhu et al. 2022). However, these findings should be interpreted cautiously since these manipulations are likely to have produced activation all the collaterals associated with these neurons and not only those projecting to the PVT. The data also present the intriguing possibility that a descending GABAergic projection from the BSTDL/CeL (Moga and Gray 1985; Ye and Veinante 2019) could dampen the excitatory effects of arousal-related signals on LPB-PVT neurons. Effective modulation of excitatory inputs could be profound if descending GABAergic inputs synapse on the soma or proximal dendrites of LPB-PVT neurons (Hao et al. 2009; Kandel and S.A. 2000). Such a multisynaptic circuit could provide a mechanism by which the BSTDL/CeL could gate ascending signals related to homeostatic state of the body to the PVT and other areas of the forebrain. Changes in this circuitry could lead to maladaptive hyperaroused states and dysregulation of homeostasis.

**Fig. 17.**
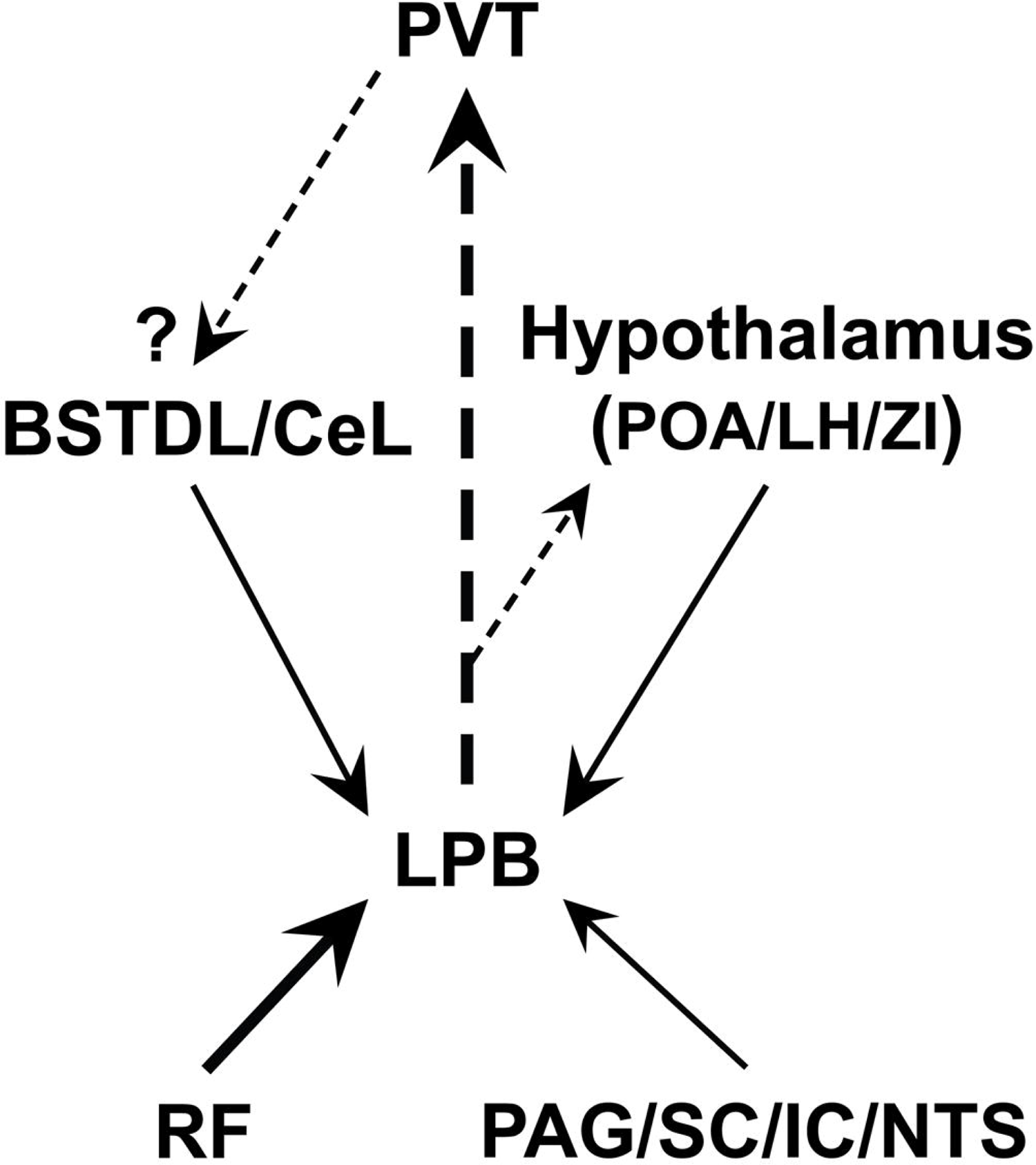
Schematic diagram summarizing some of the more salient anatomical findings from the TRIO and anterograde tracing approaches. The size of the arrows indicates the strength of the projection whereas the arrow with the dashed line indicates the lack of evidence for synaptic connectivity. See list for abbreviations.

## Abbreviations

3V: third ventricle

4V: fourth ventricle

7N: facial nucleus

ac: anterior commissure

aq: aqueduct

BLA: basolateral amygdala

BMA: basomedial amygdala

BSTDL: bed nucleus of stria terminalis, dorsolateral

BSTL: bed nucleus of stria terminalis, lateral

BSTM: bed nucleus of stria terminalis, medial

BSTV: bed nucleus of stria terminalis, ventral

BSTVL: bed nucleus of stria terminalis, ventrolateral

CeL: central amygdala, lateral-capsular division

CeM: central amygdala, medial

CnF: nucleus cuneiformis

cp: cerebral peduncle

CPu: caudate putamen

CTB: cholera toxin subunit B

DMH: dorsomedial hypothalamus

DR: dorsal raphe

dscp: decussation superior cerebellar peduncle

DT: dorsal tegmental nucleus

f: fornix

Hb: habenular nucleus

ic: internal capsule

IC: inferior colliculus

IMD: intermediodorsal nucleus of thalamus

IPAC: interstitial nucleus of the posterior limb of the anterior commissure

KF: Killiker-Fuse nucleus

LC: locus ceruleus

LH: lateral hypothalamus

LPB: lateral aspect of the parabrachial nucleus

LPO: lateral preoptic area

LRt: lateral reticular nucleus

LV: lateral ventricle

MD: mediodorsal nucleus of thalamus

MdRt: medullary reticular nucleus

Me5: mesencephalic trigeminal nucleus

mfb: medial forebrain bundle

MGN: medial geniculate nucleus

ml: medial lemniscus

MnPO: median preoptic nucleus

MPB: medial aspect of the parabrachial nucleus

MPO: medial preoptic nucleus

Mt: mammillothalamic tract

NAcSh: shell of the nucleus accumbens

NTS: nucleus of the solitary tract

opt: optic tract

PAG: periaqueductal gray

PAGD: periaqueductal gray, dorsal

PAGV: periaqueductal gray, ventral

PB: parabrachialc nucleus

PCRt: parvicellular reticular nucleus

PLH: perifornical lateral hypothalamus

POA: preoptic area

PSTh: parasubthalamic hypothalamus

PT: paratenial nucleus of the thalamus

PVN: paraventricular nucleus of the hypothalamus

PVT: paraventricular nucleus of the thalamus

aPVT: anterior aspect of the paraventricular nucleus of the thalamus

pPVT: posterior aspect of the paraventricular nucleus of the thalamus

py: pyramidal tract

RF: reticular formation

mRF: mesencephalic reticular formation

mdRF: medullary reticular formation

pRF: pontine reticular formation

rs: rubrospinal tract

SC: superior colliculus

scp: superior cerebellar peduncle

scpd: superior cerebellar peduncle, descend limb

sm: stria medullaris of the thalamus

SNL: lateral substantia nigra

sp5: spinal trigeminal tract

Sp5: spinal trigeminal nucleus

Sp5C: spinal trigeminal nucleus, caudal

STh: subthalamic hypothalamus

TH: tyrosine hydroxylase

TRIO: tracing the input-output organization method

Ve: vestibular nucleus

VLM: ventrolateral medulla

VMH: ventromedial hypothalamus

VP: ventral pallidum

ZI: zona incerta

